# Genomic basis for resistance to acute oak decline and mildew infection in English oak

**DOI:** 10.64898/2026.03.04.709495

**Authors:** Rômulo Carleial, Louise Gathercole, Gabriele Nocchi, Freya Cornwell-Davison, Sandra Denman, Nathan Brown, Richard Buggs

## Abstract

English oak (*Q. robur*) in Britain is at risk from a young age due to powdery mildew and at maturity due to acute oak decline (AOD). The contribution of oak genetic diversity to these conditions is a critical question that until recently would have required extensive clonal or family trials of juvenile and mature oak trees. Here we overcome this practical burden with genomic approaches inspired by medicine. We sequence 1868 oak trees from 78 sites and identify 1491 trees exclusively of the species *Q. robur*, which we analyse. Using single nucleotide polymorphism (SNP) based estimates of heritability (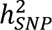), we estimate 20.9% of variation in AOD presence is genetically determined. For mildew symptoms, 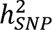 is 27.8%. Using genome-wide association studies, no SNP markers show a significant association with AOD, but 183 are significantly associated with mildew. Genomic prediction models for AOD and its symptoms, using thousands of loci, give low to moderate accuracies, ranging from 0.187 to 0.617. These results suggest a highly polygenic heritable component of susceptibility to AOD, which perhaps reflects the number of biotic and abiotic factors known to contribute to it. In contrast, powdery mildew, caused by a single fungus, has an oligogenic heritable component controlled by fewer loci of large effect. These results could be used to inform breeding programs to develop trees more resistant to AOD and mildew.

## MAIN

Oaks (*Quercus*) form a diverse genus of angiosperms (Kremer & Hipp, 2020), whose species have long been used by humans (Gil-Pelegrín et al., 2017). In Britain, the two native species, *Quercus robur* and *Q. petraea* serve as habitat and food resource for more than two thousand species (Mitchell et al., 2019), making them key species for conservation and reforestation. Both have economic importance, with *Q. robur* (English oak) as the principal broadleaf timber species in England (Gathercole et al., 2026).

In Britain, thousands of *Q. robur* are suffering from acute oak decline (AOD), a complex syndrome characterised primarily by fluid exudation from necrotic inner bark lesions (Denman & Webber, 2009; Denman et al., 2014; Brown et al., 2016; Denman et al., 2022). First formally categorised in 2009 (Denman & Webber, 2009), AOD has caused many trees to die each year and has since expanded to most areas of southern England and Wales, with a recent northern expansion, most likely driven by increases in temperature (Forestry Commission, 2023). Similar symptoms have been observed in continental Europe (Thomas et al., 2002; Thomas, 2008; Carluccio et al., 2025).

Abiotic predisposing factors associated with AOD include: lower incidence of rainfall, high temperature, high levels of nitrogen deposition, soil pH and exchangeable magnesium (Brown et al., 2018, Scarlett et al., 2021; Shaw et al., 2025). Biotic associations with AOD include the native bark beetle *Agrilus biguttatus* (Coleoptera: Buprestidae) which has seen recent population growth (Brown et al., 2015, 2017; Reed et al., 2018), and three bacteria species (*Gibbsiella quercinecans*, *Brenneria goodwinii*, *Rahnella victoriana*) which are found in necrotic tissues underlying the weeping patches in affected trees (Brady et al., 2017; Denman et al., 2018; Doonan et al., 2019, 2020). Necrosis due to bacterial infection is worsened in the presence of *Agrilus* larvae (Denman et al., 2018). While previous studies have helped elucidate the environmental and biotic risk factors associated with AOD, there is presently no information regarding its genetic risk factors. If there were a heritable component of AOD, selection of resistant germplasm could be a feasible management intervention.

Powdery mildew, mainly caused by *Erysiphe quercicola* and *E. alphitoides*, was first reported in Europe in 1907 and has since become one of the most common diseases of oak in the region (Marçais & Desprez-Loustau, 2014). It is thought to have been accidentally introduced from tropical regions, perhaps on mangos (Desprez-Loustau et al., 2017). It is not fatal to mature oaks but can contribute to their decline (Marçais & Desprez-Loustau, 2014; Sanchez Lucas et al., 2025). It has more severe effects on young trees (Marçais & Desprez-Loustau, 2014) and has been proposed as an explanation for the widespread failure of natural regeneration in *Q. robur* over the past century (Demeter et al., 2021). Some observations suggest that some natural resistance to powdery mildew may exist in native oak populations (Ayres, 1976; Lonsdale, 2015) and that the incidence and severity of powdery mildew are significantly lower on *Q. petraea* (Marçais & Desprez-Loustau, 2014; Lonsdale, 2016).

Two genomic approaches have been widely used to investigate the heritable component of disease traits. Genome-wide association studies (GWAS) seek associations between genetic markers and a phenotypic trait by estimating each marker effect independently. Abundantly used in human (Gallagher et al., 2018) and crop research (Pino Del Carpio et al., 2019; Tibbs Cortes et al., 2021), GWAS has led to the discovery of many quantitative trait loci (QTLs) underpinning disease resistance/susceptibility and commercially relevant phenotypes. It is particularly effective when traits have a genetic architecture composed of few markers of large effect. A recent GWAS study on the health of oaks under powdery mildew infection, based on 819 SNPs genotyped in 1185 individuals found 14 loci that had significant associations, mainly on chromosomes 2, 6, and 8 of the Darwin Tree of Life genome assembly (Barrès et al., 2024).

In contrast to GWAS, genomic prediction (GP) models fit very large numbers of markers simultaneously and usually assume an infinitesimal model in which each marker contributes very little to trait heritability (Meuwissen et al., 2001; Hill, 2014, Crossa et al., 2017). Models are trained using a population in which both genotypes and phenotypes are known, to estimate genomic breeding values (GEBVs) of individuals in a test population. Genomic prediction is particularly relevant in the context of germplasm selection or breeding, since individuals’ genetic merit for a trait can be estimated without the trait having been observed (e.g. in young individuals where the trait cannot yet be expressed). This can speed up breeding cycles and dramatically reduce costs (Meuwissen et al., 2001; Crossa et al., 2017; Lebedev et al., 2020). While both GWAS and GP can be used to inform selection of individuals in breeding programs, GP is more effective when traits are highly polygenic and have low heritability (Gienapp et al., 2017; McGaugh et al., 2021). Since GP models rely on linkage disequilibrium (LD) between markers and causative loci, its application for tree breeding has been considered challenging due to their large effective population sizes and low LD (Grattapaglia, 2017). To compensate for this, GP models applied to trees require very high marker densities and a large number of individuals in the training population. Due to the high costs involved, the use of GP for trees has been usually restricted to species used in very large commercial plantations, based on large progeny trials and modestly dense SNP chip markers (Grattapaglia, 2017). To our knowledge, studies aimed at predicting disease resistance in less intensively managed populations have not been widely pursued (but see Stocks et al., 2019).

Here we fill this knowledge gap by developing one of the largest genomic datasets for forest trees, composed of 1868 oak (*Quercus spp.*) whole-genomes, in order to investigate the genomic architecture of AOD and powdery mildew symptoms. Firstly, we rigorously exclude all individuals that are not diploid *Q. robur*, using analyses of population genomic structure. Secondly, we estimate SNP-heritability of AOD and mildew. Thirdly, we implement a linear mixed model GWAS to investigate the association between genetic markers and the presence of AOD and powdery mildew. Fourthly, we use genomic prediction models to generate genomic estimated breeding values and correlate these with AOD symptoms.

## RESULTS

### Sequence data and quality control

For 1996 oak samples from 77 study sites (Figure 1) we called 200,559,516 SNPs across 12 chromosomes and 538 scaffolds. Before filtering SNPs for maf > 0.05 and extreme deviations from Hardy-Weinberg equilibrium before downstream analyses, individuals had an average call rate of 0.952 ± 0.013 SD (Supplementary Figure 1) and an average coverage of 22.88x.

**Figure 1.**
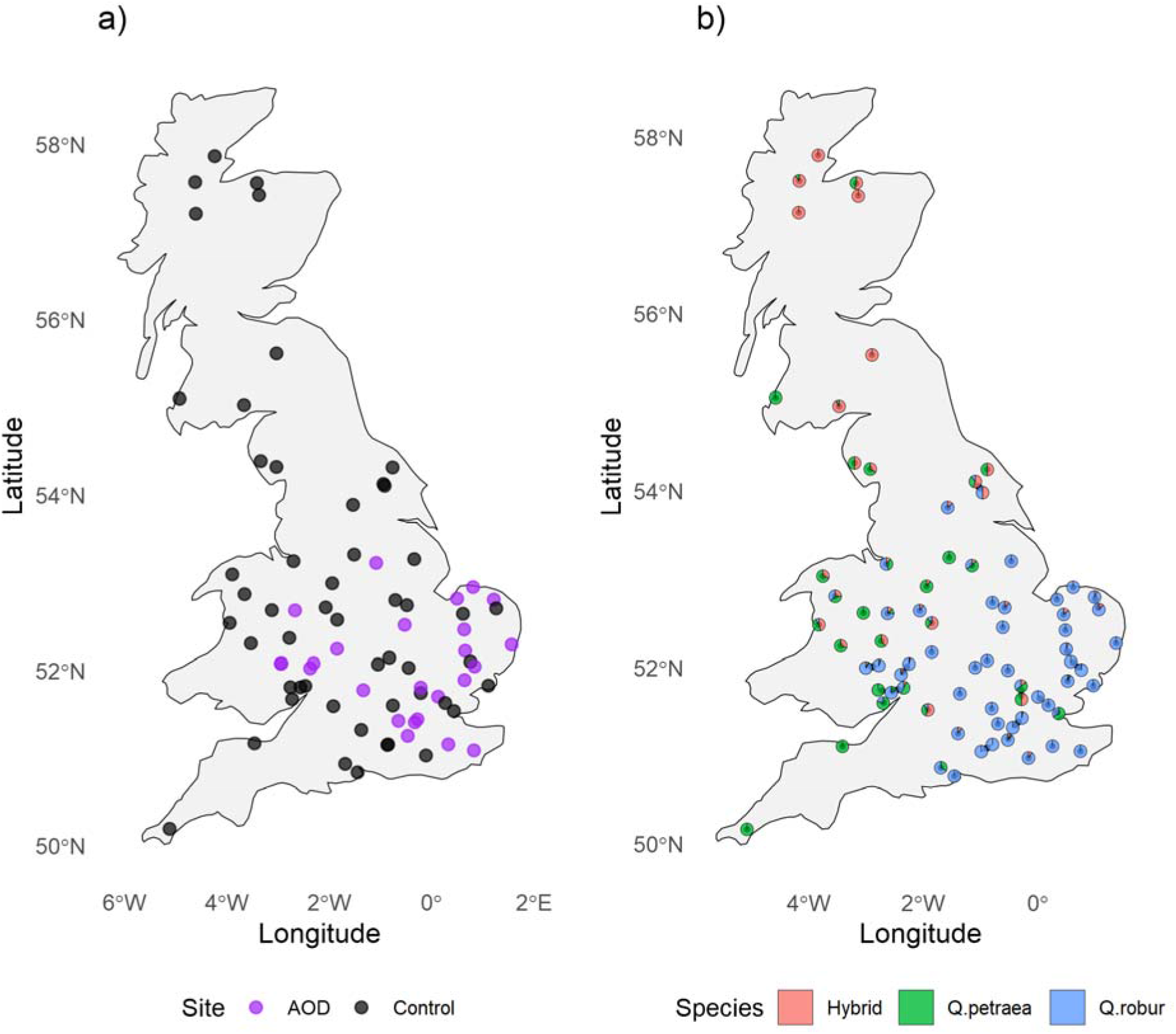
UK sites sampled for the present project (n = 77) with respect to: a) AOD occurrence and b) species prevalence. Species were inferred by admixture proportions (see Results), with any sample with over 10% admixture being designated as a hybrid.

Replicated samples were identified and each replicate with the lowest SNP call rate was removed. Samples with high missingness rates and abnormally high heterozygosity were discarded (Supplementary Figure 1) including 15 samples distributed across 11 sites that we identified as triploid (Supplementary Figure 2). The final dataset after sample quality control and SNP filtering contained 8,474,685 high-quality SNPs (**gwas snp set,** see Methods) and 1868 samples.

### Population structure analyses

The PCA plot (Figure 2a) showed two clusters and a continuum between them in PC1, which explained 61.83% of the variation (Figure 2a). These correspond to *Q. robur* and *Q. petraea* and hybrids between them. Only 6.5% of variation was summarized by PC2 (Figure 2a), where clusters were mostly caused by groups of highly related individuals (i.e. 1^st^ and 2^nd^ degree relations) from two sites, Langdale and Blickling, (Supplementary Figure 3). Admixture analyses with fastSTRUCTURE and ChooseK model comparison suggested two to four population clusters. An admixture plot assuming K=2 (Figure 2b), showed that 10.2% of the total number of individuals in our dataset were *Q. petraea*, 10.0% had 10% to 90% hybrid ancestry, and the remaining 79.8% were *Q. robur* (Supplementary Table 1). Most *Q. petraea* and hybrids were in Scotland, Wales and the west of England (Figure 1b).

**Figure 2.**
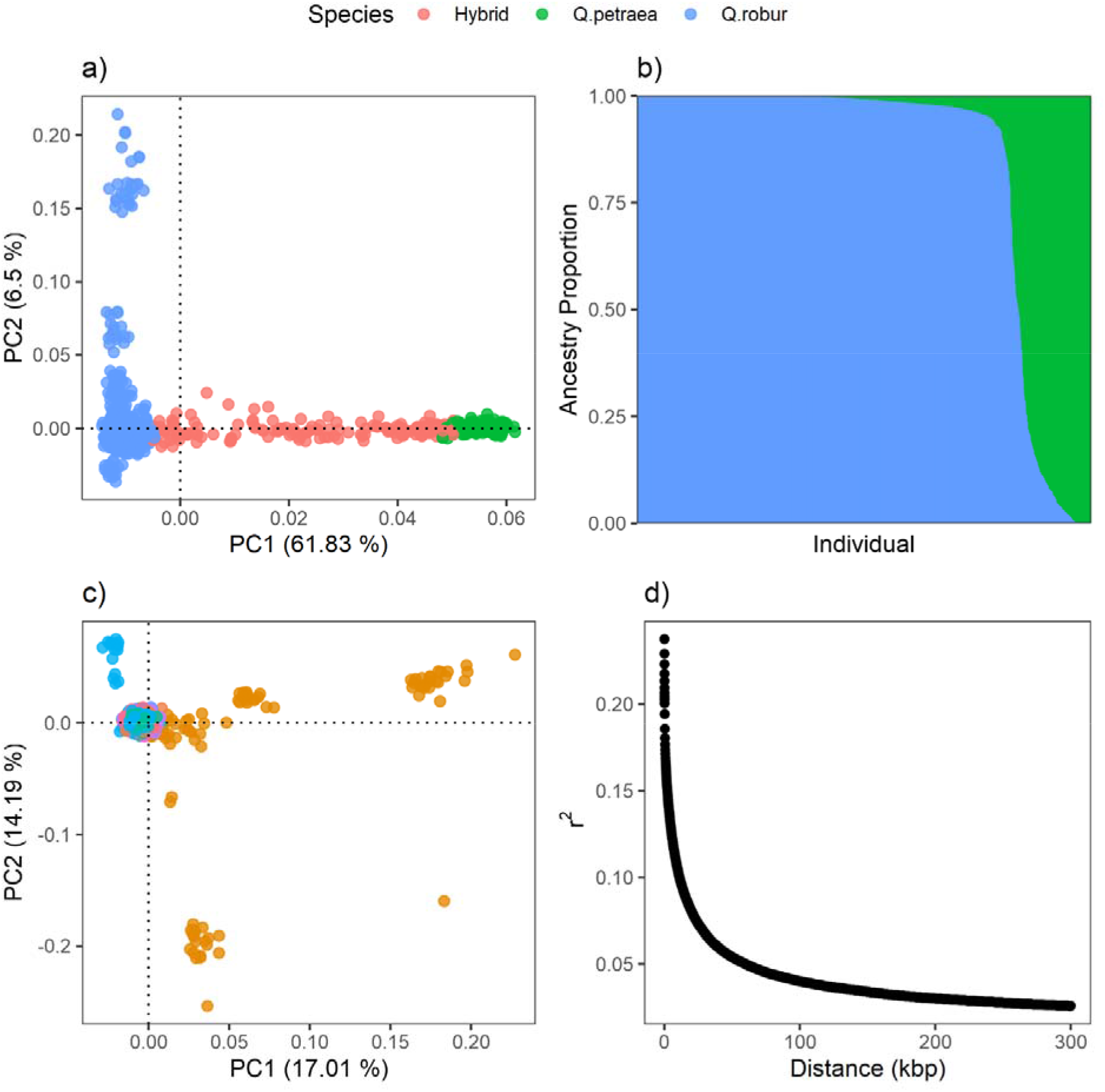
Genetic diversity of oak populations in the UK. a) Principal component analysis (PCA) using genome-wide markers and including all samples; b) admixture plot generated using a fastSTRUCTURE output assuming K=2. X-axis refer to the individual samples ordered by the degree of admixture, and y-axis the proportion of the genotype ascribed to a given species. Samples in a) and b) are colour coded by species inferred in the admixture analysis (blue = *Quercus robur*, green = *Q. petraea*, red = hybrid); c) the two main principal components summarising variation in 1440 *Q. robur* individuals. Dots are colour coded according to the collection site (n=56). Many of the outlier individuals in brown (Blickling) and light blue (Langdale) were shown to be highly related (see Results); d) whole-genome linkage disequilibrium decay in *Q .robur*.

Retaining the 1491 *Q. robur* individuals we checked for any population substructure associated with geographic distribution which could confound GWAS. In PCA, PC1 and PC2 separated the above mentioned Langdale and Blickling outliers, but there was no evidence for strong population substructure across the remaining sites (Figure 2c, Supplementary Figure 4). PCAs including only unrelated or distantly related individuals (i.e. less than 3^rd^ degree relatives) no longer separated Langdale from the other sites, but Blickling still showed signs of differentiation (Supplementary Figure 5) perhaps due to planting. In pairwise F_st_ comparisons, we found very little differentiation between populations (mean F_st_ = 0.007), with Coed Penrhyn in the westernmost part of Wales and Maldon Wood in the easternmost part of England being the most differentiated (F_st_ = 0.052). We found whole-genome linkage disequilibrium to decay in *Q. robur* at very short distances. Correlation coefficients between SNPs usually dropped below 0.2 (i.e. r^2^ < 0.2) after 99 bp and below 0.1 after 11,205 bp (Figure 2d) suggesting high recombination rates.

### SNP heritability

We estimated the SNP heritability (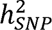) of phenotypic traits in *Q. robur* using LDAK v6 (Speed et al., 2017) with a restricted maximum likelihood (REML) regression. For AOD status (0=control, 1=case) we found heritability of 0.209 (s.d. = 0.142). For presence of *Agrilus* exit holes (0=control, 1=case) we found heritability of 0.157 (s.d. = 0.141), for canopy absolute transparency 0.157 (s.d. = 0.192), and for stem bleeds (discrete, with both active and inactive bleeds included) 0.075 (s.d. = 0.153). For presence of mildew symptoms (0=control, 1=case) heritability was 0.278 (s.d. = 0.234) (Table 1), a higher value than we found for any AOD associated symptoms.

**Table 1.** SNP heritability estimates for mildew and acute oak decline and its symptoms.

| Phenotype | n | Prevalence* | SNP-H2 | LRT | p | Correlation | Accuracy |
| --- | --- | --- | --- | --- | --- | --- | --- |
| Acute oak decline | 1096 | 0.3914 | 0.2090.142 | 2.319 | <0.001 | 0.1030.017 | 0.225 |
| *Agrilus* exit holes | 1089 | 0.1451 | 0.1570.141 | 1.322 | <0.001 | 0.1190.016 | 0.3 |
| Canopy absolute transparency | 793 | - | 0.1570.192 | 0.630 | <0.001 | 0.0740.019 | 0.187 |
| Mildew | 657 | 0.0943 | 0.2780.234 | 1.427 | <0.001 | - | - |
| Stem bleeds | 1000 | - | 0.0750.153 | 0.238 | <0.001 | 0.1690.023 | 0.617 |
| Notes: = SNP heritability; LTR = likelihood ratio test; * percentage of cases among samples; Correlation = Pearson's correlation between genomic estimated breeding values and phenotypes; Accuracy = correlation |  |  |  |  |  |  |  |

**Table 2.** Within-gene SNPs significantly associated with biotic stressors in pedunculate oak (*Quercus robur*.

| Phenotype | Chr | SNP (bp) | -log<sub>10</sub>(p) | MAF | Gene ID | Putative function |
| --- | --- | --- | --- | --- | --- | --- |
| *Agrilus* exit holes | 3 | 1255063 | 7.265 | 0.094 | Qrob_H2.3_Sc0000136 | Inorganic pyrophosphatase |
| Mildew | 2 | 58412231 | 6.394 | 0.113 | Qrob_H2.3_Sc0000087 | Peptide-methionine (R)-S-oxide reductase |
| Mildew | 2 | 58412940 | 6.049 | 0.111 | Qrob_H2.3_Sc0000087 | Peptide-methionine (R)-S-oxide reductase |
| Mildew | 2 | 58413317 | 6.414 | 0.113 | Qrob_H2.3_Sc0000087 | Peptide-methionine (R)-S-oxide reductase |
| Mildew | 3 | 10615801 | 6.594 | 0.240 | Qrob_H2.3_Sc0000189 | Dosage compensation complex subunit MLE // Copper chaperone |
| Mildew | 6 | 18856343 | 6.888 | 0.065 | Qrob_H2.3_Sc0000102 | Ammonium transporter |
| Mildew | 6 | 25097911 | 6.137 | 0.159 | Qrob_H2.3_Sc0000196 | Histone H3 (Lys9) methyltransferase SUV39H1/Clr4 |
| Mildew | 7 | 41400821 | 6.014 | 0.062 | Qrob_H2.3_Sc0000101 | Heme-binding protein-related |
| Mildew | 7 | 41400928 | 5.972 | 0.068 | Qrob_H2.3_Sc0000101 | Heme-binding protein-related |
| **Mildew** | 7 | 49125549 | 7.038 | 0.086 | Qrob_H2.3_Sc0000058 | Phosphatidylinositol N-acetylglucosaminyltransferase subunit P-like protein |
| **Mildew** | 11 | 19058271 | 5.980 | 0.123 | Qrob_H2.3_Sc0000057 | Alcohol dehydrogenase class-P |
| **Mildew** | 11 | 19120896 | 6.492 | 0.097 | Qrob_H2.3_Sc0000057 | Alcohol dehydrogenase |
| **Mildew** | 11 | 19265181 | 6.203 | 0.127 | Qrob_H2.3_Sc0000057 | L-galactose 1-dehydrogenase |
| **Mildew** | 11 | 19265648 | 6.338 | 0.127 | Qrob_H2.3_Sc0000057 | L-galactose 1-dehydrogenase |
| **Mildew** | 11 | 19266418 | 6.675 | 0.128 | Qrob_H2.3_Sc0000057 | L-galactose 1-dehydrogenase |
| **Mildew** | 11 | 19343455 | 6.104 | 0.093 | Qrob_H2.3_Sc0000057 | Leucine Rich Repeat |
| **Mildew** | 11 | 19344854 | 6.264 | 0.092 | Qrob_H2.3_Sc0000057 | Leucine Rich Repeat |
| **Mildew** | 11 | 19378871 | 6.016 | 0.136 | Qrob_H2.3_Sc0000057 | Domain of unknown function (DUF313) |
| **Mildew** | 11 | 19405382 | 6.549 | 0.131 | Qrob_H2.3_Sc0000057 | Glutamate-1-semialdehyde 2,1-aminomutase |
| **Mildew** | 11 | 19408182 | 6.278 | 0.132 | Qrob_H2.3_Sc0000057 | Glutamate-1-semialdehyde 2,1-aminomutase |
| **Mildew** | 11 | 19410139 | 6.066 | 0.130 | Qrob_H2.3_Sc0000057 | Glutamate-1-semialdehyde 2,1-aminomutase |
| Note: bp = base pair; Chr = chromosome; MAF = minor allele frequency. |  |  |  |  |  |  |

### Genome-wide association study (GWAS)

We tested for associations between genetic markers and phenotypes in *Q. robur* using a linear mixed model (LMM) implemented in LDAK v6 (Speed et al., 2017) with site environmental variables as covariates (Supplementary Table 2). We found no significant associations between SNPs and AOD status irrespective of the p-value threshold used, suggesting that the heritable component of variation in AOD resistance is modulated by many genes of small effect in *Q. robur*. Nevertheless, the Manhattan plot showed some clear peaks just below the significant threshold, which could suggest we are lacking in statistical power (Figure 3; Supplementary Figure 6). A list of putative genes and their genomic distance from SNPs within peaks using an arbitrary cutoff of - log10 = 5 is summarised in the Supplementary Table 3. Some of these have functions that can be linked to insect, bacterial or stress response. Two of these genes have been previously identified in genomic regions highly differentiated between *Q. petraea* and *Q. robur* (Gathercole et al., 2026).

**Figure 3.**
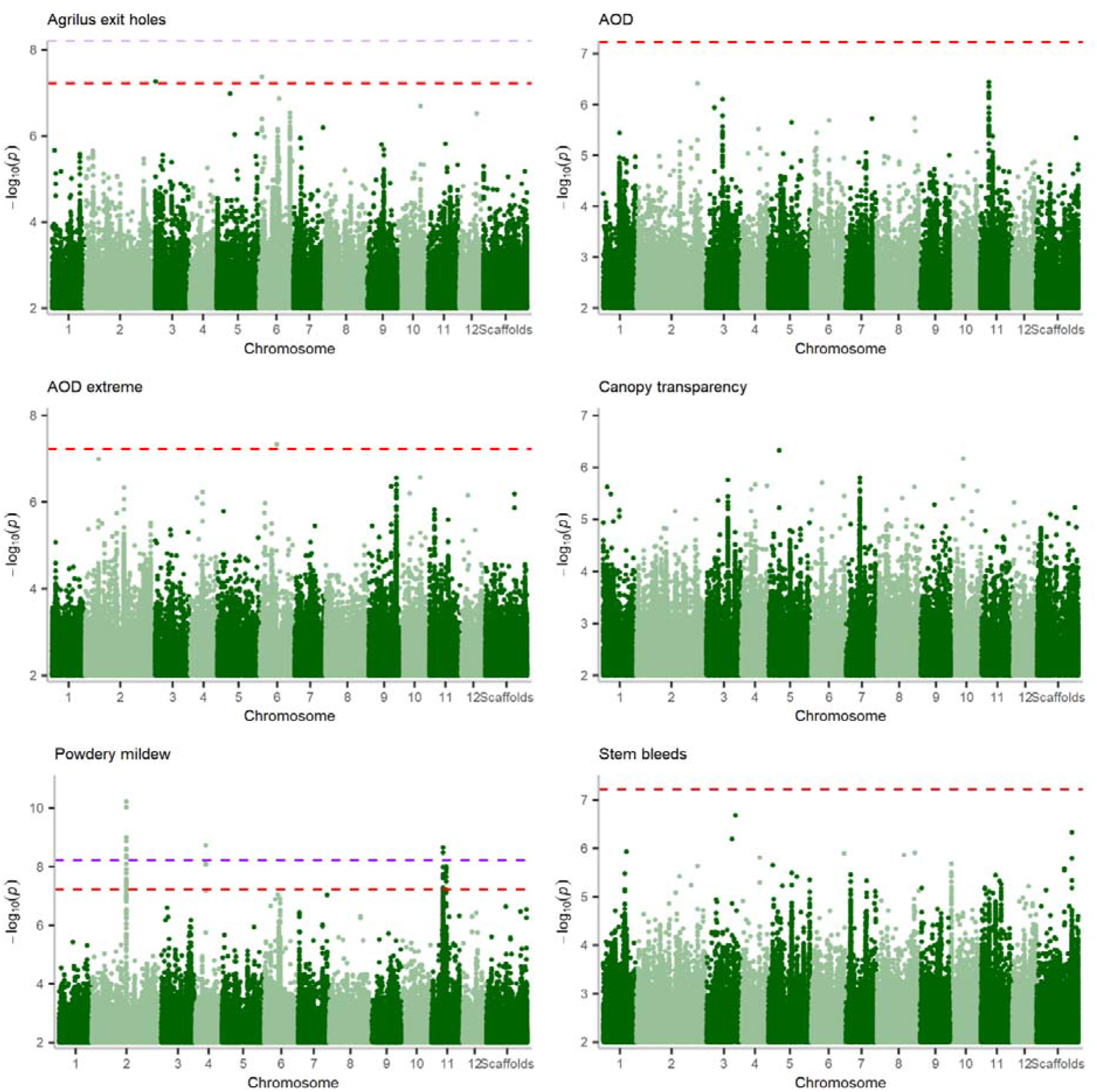
Genome-wide associations between SNPs and 6 traits of interest in pedunculate oak (*Quercus robur*). Dashed purple and red lines represent the Bonferroni and modified Bonferroni significance thresholds, respectively.

A GWAS restricted to *Q. robur* individuals at the extreme of the symptom severity spectrum i.e. either never manifested AOD symptoms for the whole duration of the monitoring period (i.e. 14 years) or had symptoms for more than half of those years, revealed one SNP which was significantly associated with AOD if a modified Bonferroni correction threshold was used (Figure 3). The SNP is located in chromosome 6 and is 752bp away from a gene annotated as large subunit ribosomal protein L25. We found two SNPs associated with presence of *Agrilus* exit holes if a modified Bonferroni threshold was used (Figure 3). A SNP in chromosome 3 is inside the coding region of a gene annotated as inorganic pyrophosphatase (Table 1). One in chromosome 6 is 3,122bp away from a gene annotated as a carboxymethylenebutenolidase homolog. No SNPs were significantly associated with the number of stem bleeds or canopy crown transparency ( Figure 3).

For crown mildew, GWAS found 183 SNPs which were significantly associated with symptoms in *Q. robur* when we used a 5% FDR significant threshold. This number reduced to 11 if the more conservative Bonferroni threshold was used ( Figure 3). It is worth noting that we found signs of genomic inflation in diagnostic QQ plots, suggesting some population structure not fully accounted for in the model (Supplementary Fig. 6). A full list of statistically significant SNPs is provided in the Supplementary Table 4. These SNPs are near (<60kbp) 44 annotated genes, five of which have been previously identified in regions of high differentiation between *Q. robur* and *Q. petraea* (Gathercole et al., 2026). Below we only report SNPs within gene coding regions, many of which are related to oxidative stress (Table 1).

In chromosome 2, three SNPs supported a gene annotated as peptide-methionine (R)-S-oxide reductase, which is involved in repairing damage from oxidative stress in plants (Rey & Tarrago, 2018). In chromosome 3, one SNP was inside the coding region of a gene annotated as a dosage compensation complex subunit MLE (involved in transcription) and copper chaperone (involved in inorganic ion transport and metabolism). Copper chaperones prevent oxidative stress and have been previously associated with immune response against bacteria in *Arabidopsis* (Chai et al., 2020). In chromosome 6, one SNP was inside the coding region of an ammonium transporter gene, which may have a role in immunity against stem rust in wheat (Li et al., 2017), and another SNP was inside the coding region of a Histone H3 (Lys9) methyltransferase gene. Similarly, a recent study found a SNP within the coding region of a histone H4 gene associated with survival in a young *Q. robur* cohort naturally exposed to mildew (Barrès et al., 2024). In chromosome 7, three SNPs were located inside the coding region of a gene. Two SNPs were inside a gene annotated as heme-binding protein-related, and another inside a phosphatidylinositol N-acetyglucosaminlytransferase subunit P-like protein gene. The latter has been associated with growth under stress conditions (Yoo et al., 2021). In chromosome 11, two SNPs were inside an alcohol dehydrogenase gene, known to be associated with abiotic stress (e.g. Shen et al., 2021), and three were inside a L-galactose 1-dehydrogenase gene, which is involved in vitamin C biosynthesis (Vargas et al., 2022). Two SNPs were inside a leucine rich repeat gene, known to be involved in plant immune response against pathogens (Padmanabhan et al., 2009), and another three were inside a Glutamate-1-semialdehyde 2,1-aminomutase gene, involved in assimilation and re-assimilation of ammonia (Unno et al., 2006). Finally, one SNP was inside a Domain of unknown function (DUF313), which has been associated with drought and salinity stress responses in rice (Guo et al., 2016).

### Genomic prediction

To test whether we could accurately predict phenotypic traits using genetic markers, we built elastic net prediction models (Hof & Speed, 2025) using LDAK v6, randomly distributing samples into a training set containing 90% of samples and a test set with the remaining 10%, repeating the procedure ten times until all samples were included in both sets. Correlations between phenotypes and genomic estimated breeding values (GEBVs) for AOD status, stem bleeds and canopy transparency were very low, ranging from 0.074 to 0.169 (Table 1). The prediction accuracy of presence of *Agrilus* exit holes was 0.3, of AOD status was 0.225, of canopy absolute transparency was 0.187 and of stem bleeds was 0.617.

For AOD, we explored the possibility that models trained with GWAS markers could increase genomic prediction accuracy. Models were run with one randomised set (10% of individuals). We obtained slightly higher genomic prediction accuracies (r^2^ = 0.208 ± 0.02 SD) when we trained ENET models with the top 10k GWAS predictors (Figure 4), than including all markers. The inclusion of more GWAS markers decreased accuracy progressively. Predictions trained with markers picked at random had lower accuracy than GWAS markers if 10k or less markers were used, but were more accurate at 500k markers. Models trained with all genomic markers (n = 8,474,685) had similar accuracy (r^2^ = 0.194 ± 0.026 SD) than models trained using 500k random markers (r^2^ = 0.194 ± 0.012 SD), suggesting increasing marker density becomes uninformative after a certain point.

**Figure 4.**
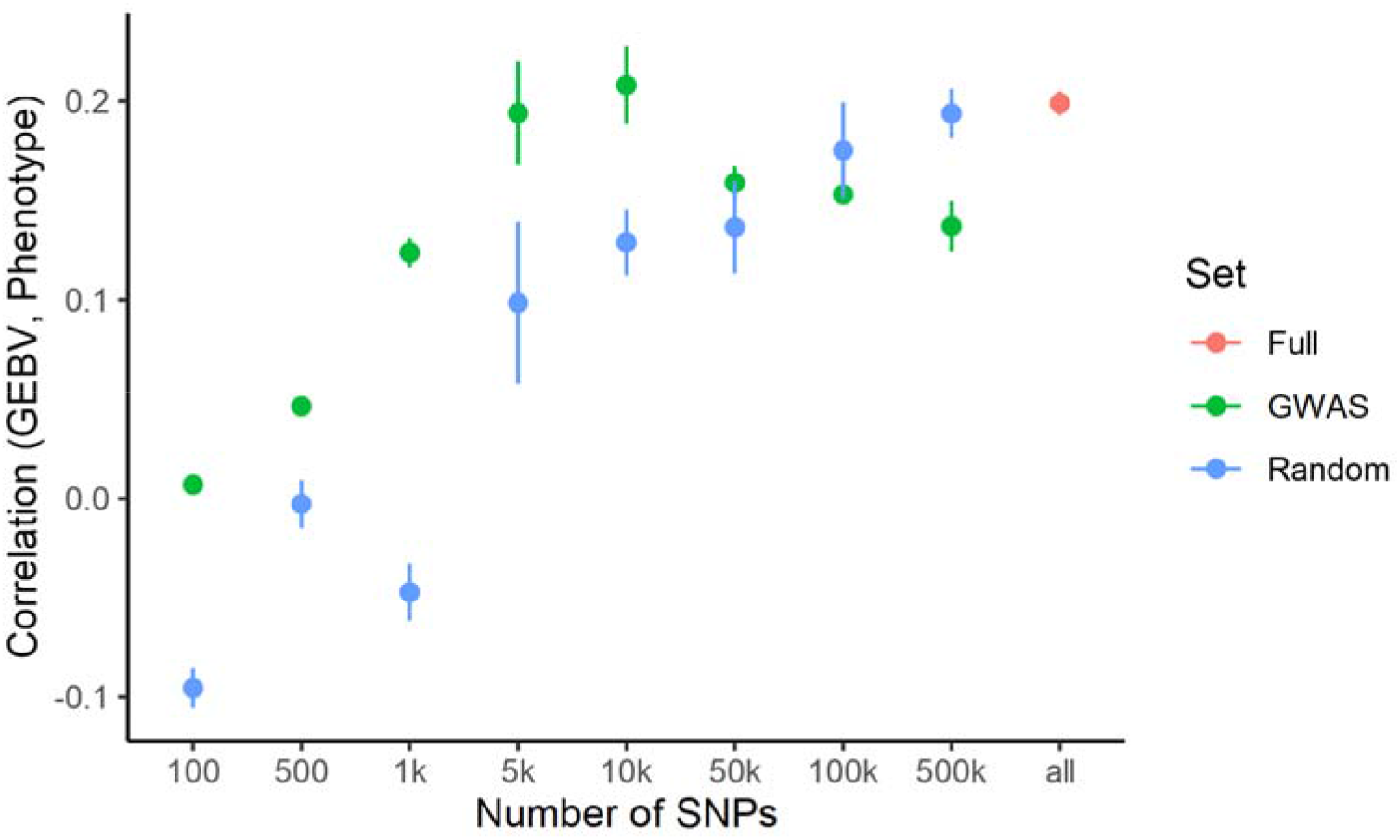
Impact of the number and origin of SNPs in the accuracy of genomic prediction for acute oak decline in pedunculate oak (*Quercus robur*).

## DISCUSSION

Collectively, these results are a step forward in our understanding of AOD and mildew susceptibility in English oak. Our estimate of 0.21 heritability for AOD and 0.28 heritability for powdery mildew means that breeding for reduced symptoms could be feasible using genomic selection. In livestock, traits with lower levels of heritability than this have been improved by genomic selection (Samorè & Fontanesi, 2016; Tade & Melesse, 2024). In loblolly pine, stem volume and stem straightness have estimated heritability of 0.11 to 0.22 and are amenable to improvement by genomic selection (Isik et al., 2026). In interpreting our heritability estimated for oak, it should be borne in mind that the trees in our study are not growing in uniform and controlled environments. We would expect heritability to be lower in woodlands and parks than it would be if traits were measured in a planted trial, where trees were uniform in age and spacing, and where weeds and herbivores are controlled. Our heritability estimate for mildew was at the lower end of the range of genomic narrow-sense heritability (h² = 0.25–0.67) of mildew score estimated within a *Q. robur* F_1_ mapping family across two field sites and multiple years (Bartholomé et al., 2020). Many studies have shown relationships between concentrations of nutrients present in the soil and susceptibility to AOD (Brown et al., 2018, Scarlett et al., 2021; Shaw et al., 2025), and such concentrations vary both within and between sites. The symptoms of AOD can also differ seasonally and between years, so the fact that some of our trees were phenotyped in different months and years may have also reduced our heritability estimates. The high variation in AOD and mildew attributable to environment in our estimates of heritability suggest that immediate management interventions to modify micro-environments, perhaps by enriching the soil of badly affected parklands and woodlands, could reduce the incidence or severity of both problems.

From our sequence data from 1491 *Q. robur* individuals, despite some clear peaks in the GWAS Manhattan plots, we did not find individual single nucleotide polymorphisms (SNPs) significantly associated with AOD, suggesting that heritable resistance is polygenic. Attempting to find associations for individual symptoms independently, or restricting the analysis to individuals at the more extreme ends of the susceptibility spectrum, gave similar results. A better understanding of the causal factors behind AOD will in principle lead to more fine-grained phenotyping (Finch et al., 2021), which should increase the statistical power to detect associations in the future.

Nevertheless, examining the highest peaks in our Manhattan plots reveals loci with potential involvement in biotic or abiotic stress resistance. For example, a peak in chromosome 1 contains SNPs which are close to a gene which has a homolog in *Arabidopsis* which is associated with aluminium tolerance. Anecdotal evidence from Britain has suggested soil aluminium is associated with AOD (see report for Defra FPPH Project PH0469, 13/05/2020), perhaps because soil aluminium reduces water and nutrient uptake. However, given that soil aluminium concentrations are site specific, not all trees in our dataset would have been exposed to aluminium stress. To confirm this finding, we would need to run a GWAS that only includes sites with certain aluminium levels, but running a GWAS per site would be unfeasible with our current data, as it would reduce the sample size and statistical power by a large margin. It is therefore possible that some GWAS peaks we identified may be indicative of true causal variants whose effects are partially masked by environmental variability. Future studies would benefit from sampling individuals in the same year and within the span of a few weeks, which should reduce environmental noise. A more experimental solution would be to develop screening trials in a common environment (e.g. Plumb et al., 2019), which should also reduce the confounding effects of woodland age and surveyor bias, known to affect the scoring of a tree health status (Zarnoch et al., 2004).

In contrast to AOD, for powdery mildew symptoms we identify markers with larger effects, many of which are related to oxidative stress. Oxidative stress has been previously linked to cell death as a reaction to powdery mildew in wheat (Li et al., 2019). A recent study with *Q. robur* half-sib families in controlled experiments with treatments protected or not protected against mildew infection, identified three candidate genes potentially involved in mildew susceptibility and survival (Barrès et al., 2024). One of these genes, a pentatricopeptide repeat-containing protein (PPR) gene located in chromosome 2, was also identified in our study although in a different chromosome (i.e. chromosome 6, Supplementary Table 4). It is important to notice however, that the investigators used a different *Q. robur* reference genome (Lang et al., 2021), and that this particular gene was identified in a GWAS including only those individuals protected against mildew infection by fungicide application (Barrès et al., 2024). In the unprotected treatment, Barrès et al. (2024) found a putative histone H4 gene in chromosome 8 associated with survival in the mildew exposed treatment, while we found a putative histone H3 gene in chromosome 6. Our combined results suggest the roles of histones and PPR genes in mildew susceptibility warrant further investigation.

We compared SNPs and nearby genes identified as associated with AOD and mildew in *Q. robur* with those previously identified as occurring in genomic regions of high differentiation between *Q. robur* and *Q. petraea* (Gathercole et al., 2026). In high differentiation regions we found four genes exclusively associated with mildew, one exclusively associated with AOD, and one common to both. However, most of our candidate genes occurred in regions where *Q. robur* and *Q. petraea* seem to be exchanging alleles freely. Despite this, we found that 94.54% of the 183 SNP alleles significantly associated with mildew occur at higher frequencies in *Q. robur* (MAF = 0.14 +-0.103) than in *Q. petraea* (MAF = 0.045 +-0.124). Previous studies have shown that the incidence and severity of powdery mildew are significantly higher on *Q. robur* than on *Q. petraea* (Marçais & Desprez-Loustau, 2014; Lonsdale, 2016), so our results suggest that this could have a genetic component. Hybridisation between species can be a source of genetic variation allowing rapid adaptation to new conditions (Petit et al., 2004; Edelman & Mallet, 2021). Our results hint that introgression of alleles from *Q. petraea* could in the future help *Q. robur* to develop greater resistance to powdery mildew. Currently there has not been time for this to happen as powdery mildew is a recent invader relative to the age of the trees included in this study. In contrast to mildew, we found that only 27.87% of the alleles at 77 loci related to AOD symptoms are more frequent in *Q. robur*, which might suggest that *Q. petraea* is more susceptible. This suggestion should be treated with great caution as a highly polygenic trait affected by many loci of small effect is very hard to predict in different genetic backgrounds. There is presently no systematic evaluation of whether *Q. robur* is more susceptible to AOD than *Q. petraea*. The higher incidence of AOD in British *Q. robur* populations has been usually attributed to its southern distribution in the UK, where AOD risk factors are more prevalent (Brown et al., 2018).

We were able to predict AOD symptoms with low to moderate accuracy. Similarly, a recent study attempted to predict the phenotypes of unrelated plus trees and generally found low accuracy for most traits (Hiraoka et al., 2018). Polygenic traits with low heritability are suitable for genomic prediction (Crossa et al., 2017; Grattapaglia, 2022), and models have been successfully applied in the context of crop or tree breeding for higher yield or timber quality (Crossa et al., 2017; Grattapaglia, 2022; Isik et al., 2026). Thus far, GP models have been little used for breeding disease resistance in crops and trees (Poland & Rutkoski, 2016; Lebedev et al., 2020), but selecting for AOD resistant trees, even when predictions have low accuracy, should still provide genetic gains and be advantageous in restoration initiatives or forestry provided other better characterised traits are not compromised during selection. Our results suggest that using the number of stem bleeds as a proxy for AOD leads to higher prediction accuracy, so it should be the trait of choice if our models are to be used for tree selection. However, genomic selection programs for AOD resistance based on our findings would likely require that selected trees be somewhat related to the trees used here. GP models have been shown to lose at least 20% accuracy when the training and test populations are unrelated (Beaulieu et al., 2014; Müller et al., 2017; Carleial et al., 2025).

In conclusion, our results support the hypothesis that AOD resistance is a polygenic trait, while mildew resistance seems to be driven by fewer loci of larger effect. Management strategies for AOD focusing on managing the tree micro-environment, particularly soil composition and water availability may therefore be more appropriate. Nevertheless, our genomic prediction models for AOD performed acceptably given the low heritability, and provide moderate accuracies, meaning that genomic selection for reduced AOD susceptibility is possible. For mildew, marker assisted selection would be the most promising approach to increase oak population resilience to this threat.

## METHODS

### Phenotyping and sample collection

Oak samples used in this study were collected in four tranches including sites spanning most of Britain (Figure 1). The first tranche, collected in 2017, included 458 samples from four sites and has been described extensively in Nocchi et al. (2021). These samples have been previously analysed in terms of population structure (Nocchi et al., 2021), and the microbiome (Gathercole et al., 2021). A second tranche was collected in the summer of 2019 by the Forest Research Technical Services Unit, of which 424 samples and 15 technical replicates from 60 Forest Condition Survey sites were successfully sequenced. This tranche was previously analysed with respect to population structure, hybridisation and ploidy (Gatherchole et al., 2026). The third tranche was collected in 2021 with the addition of 700 samples from 24 sites, 22 of which were novel. In 2022, a fourth tranche consisting of 400 samples was added from four novel sites. Most samples in tranches 1, 3 and 4 were collected from sites which were AOD-affected, but tranche 2 was Britain-wide with no AOD requirement in the site selection (Figure 1a) (Gathercole et al., 2026). The sampling strategy for AOD-affected sites in tranches 1, 3 and 4 involved collecting data from healthy and symptomatic trees in equal proportions, which invariably led to a dataset with a higher proportion of symptomatic trees than in the wider British population (i.e. ascertainment bias). Trees were classified as either AOD-affected or healthy by well-trained surveyors during sample collection. Classification was based on a minimum set of AOD symptoms which included canopy absolute transparency, presence or absence of *Agrilus* exit holes, and presence or absence of stem bleeds (active or inactive). Information on the presence or absence of mildew infection on leaves was also collected. Due to time constraints during sampling, many trees in tranches 1 and 3 were limited to these phenotypes.

For tranches 2 and 4, and some trees in tranche 3, we were able to collect fine-grained measures of stem bleeds, *Agrilus* exit holes and mildew. Leaf samples were taken from each tree and dried in plastic bags containing silica gel. These were sent to the Jodrell Laboratory at the Royal Botanic Gardens, Kew for collation and storage. When possible, we collected swabs samples taken from weeping wounds which were sent to Forest Research laboratories at Alice Holt to confirm the presence of AOD-associated bacteria using a real time PCR assay technique (Crampton et al., 2020) and assist the field diagnosis. Trees which were classified as symptomatic in the field, but whose wounds did not contain AOD bacteria, were reclassified as asymptomatic.

Reclassification was restricted to trees for which wound samples were active and fresh, as old and dried samples may no longer contain pathogenic bacteria. For sites with long-term monitoring data (n = 5) (Brown et al., 2016), we were able to identify trees which were in remission i.e. had shown AOD symptoms in previous years but were asymptomatic at the time of sampling. Classifying trees in remission as either symptomatic or asymptomatic is challenging (Finch et al., 2021), but we opted for the latter to ensure consistency between sites with and without historical data.

### Sample preparation and DNA extraction

#### Tranche 1

Sequencing information for tranche one has been described in Nocchi et al. (2021). Reads were downloaded from the European Nuclear Archive (accession code: PRJEB30573).

#### Tranche 2

Sequencing information for tranche 2 has been extensively described in Gathercole et al. (2026). Reads were provided by the author (ENA accession code: PRJEB43909).

#### Tranche 3

DNA extraction, library preparation and sequencing were performed by Novogene. DNA was extracted from leaf samples and sequencing libraries were generated using NEBNext® DNA Library Prep Kit following manufacturer’s recommendations with indices added to each sample. The genomic DNA was randomly fragmented to a size of 350bp by shearing. DNA fragments were end-polished, A-tailed, and ligated with the NEBNext adapter for Illumina sequencing. The fragments with adapters were PCR amplified, size selected, and purified (AMPure XP system) and the resulting libraries were analysed for size distribution by AATI Bioanalyzer and quantified using real-time PCR. The libraries were sequenced on an Illumina NovaSeq 6000 platform with 2 × 150 bp reads. For quality control, paired reads that (1) contained adapter contamination, (2) uncertain nucleotides (N) constituted more than 10 percent of either read, (3) low quality nucleotides (base quality less than 5, Q<=5) constituted more than 50 percent of either read, were discarded.

#### Tranche 4

DNA extraction, library preparation and sequencing were performed by Edinburgh Genetics. Library Preparation was performed using *egSEQ Enzymatic DNA Library Preparation*. DNA was fragmented and selected to roughly 300-400bp insert size, followed by end repair, A-tailing and adaptor ligation. DNA was then purified and PCR amplified, followed by further purification, and circularisation pre-sequencing for DNBSEQ. Sequencing was performed on MGI DNBseq-T7 with read length PE150. Data QC and filtering was performed using soapnuke software (ref) with the following conditions: adapters were cut if the sequencing read matched 25.0% or more of the adapter sequence (maximum 2 base mismatches were allowed); sequencing reads were discarded if (1) their length was less than 150 bp, (2) the N content in the sequencing read accounted for 1.0% or more of the entire read, (3) if the length of polyX (X can be A, T, G or C) in the sequencing read exceeded 50bp, or (4) if the bases with a quality value of less than 20 in the sequencing read accounted for 30.0% or more of the entire read. The output read quality value system was set to Phred+33.

### Sequence alignment and variant calling

We quality checked the raw reads using fastqc v.0.11.9 (Andrews, 2010), which were then aligned to the *Q. robur* reference genome (Plomion et al., 2018) using bwa v.0.7.17 (Li & Durbin, 2009) with the -M option. Aligned reads were filtered with samtools v1.10 (Li et al., 2009) to remove those with mapping quality below 20 and PCR duplicates. We called variants for the 1996 samples using GATK v4.2.6.1 (McKenna et al., 2010; DePristo et al., 2011), which were then joint called following the GATK pipeline. This generated a combined VCF file containing 244,386,895 variants (200,559,516 SNPs + 43,827,379 Indels) across 12 chromosomes and 538 scaffolds. After discarding indels, we hard-filtered the remaining SNPs to eliminate low quality variants using a 4 step approach: **1)** we used vcf tools (Danecek et al., 2011) with the following options --max-missing 0.5, --minQ 30, --mac 3, --minDP 3, --min-alleles 2, --max-alleles 2 (96,393,272); **2)** we removed SNPs with very low allele frequencies with vcftools option --maf 0.01 (40,629,819 variants); **3)** we used GATK to filter the remaining SNPs with the following options: QD < 5, SOR > 3.0, FS > 10.0, MQ < 55.0, MQRankSum < -3.0, ReadPosRankSum < -2. (23,131,048 variants), and finally **5)** plink2 v2.0-20220603 (Chang et al., 2015) was used to remove SNPs with minimum allele frequency < 0.05, missing > 5% of data, and greatly deviating from Hardy-Weinberg Equilibrium (p < 1^-50^) as per recent recommendations (Reed et al., 2015; Marees et al., 2018). This led to a reduced file containing 8,474,685 high quality SNPs, which will be referred to as the **gwas SNP set**. To identify replicate samples and discover any potential mislabelling, we built a genomic relatedness matrix using plink2 --make-king function. Kinship analyses revealed most technical replicates (n = 86) matched their counterpart samples and highlighted that some trees were resampled accidentally across different project tranches (n = 27). Identities for these samples were confirmed by GPS coordinates. For subsequent analyses only the replicate sample with the highest SNP call rate was retained. Samples that came out as replicates (kinship coefficient > 0.354) but did not have matching ids (n = 18) and samples with the same id, but which were too genetically dissimilar (n = 12), were discarded as labelling errors. Moreover, since triploid individuals have been previously identified in a subset of this dataset (Gathercole et al., 2026), we plotted allele balance of samples with abnormally high heterozygosity rates using vcfR v1.15.0 (Knaus & Grünwald, 2017) and 50k randomly selected SNPs. Heterozygosity rates were calculated in LDAK v6 (Speed et al., 2017). Following Gathercole et al. (2026), individuals we identified as having many loci with an allele balance ratio of 0.33/0.67 were categorised as triploids (Fletcher et al., 2022). Twenty-eight samples had high missing rates (> 10%) and consequently low heterozygosity rates, while 27 showed abnormally high heterozygosity rates (Supplementary Figure 1). While the former are definitely poor quality samples, the latter could either be cross-contaminated samples or individuals with abnormal ploidy. Accordingly, five of these 27 individuals have been previously identified as triploids (Gathercole et al., 2026), and of the remaining 23 individuals, 10 showed unambiguous triploid patterns (Supplementary Figure 2). The confirmed triploids (n = 15) represent ∼0.82% of the evaluated samples (n = 1833) and were distributed across 11 sites. For GWAS and GP, individuals deviating more than two standard deviations from the mean heterozygosity rate were excluded, irrespective of whether they had been diagnosed as triploids or not.

Specifically for population structure analyses, we removed SNPs located in unassembled scaffolds and pruned the remaining SNPs for linkage disequilibrium (r^2^ > 0.2) using plink2 --indep-pairwise 50 5 0.2, which generated a **reduced SNP set** of 1,545,541 SNPs. Individuals which failed quality control, triploids, and replicate samples were excluded from consideration.

### Population structure analyses

The UK is home to two native (*Q. robur* and *Q. petraea*) and at least three introduced (*Q. ilex*, *Q. cerris*, *Q. rubra*) oak species. Misidentifications may occur due to morphological similarities between species, and widespread hybridisation between different species of oak have often been reported (e.g. Jensen et al., 2009; Beatty et al., 2016; Leroy et al., 2020). Accordingly, previous research with a subset of this dataset identified abundant signs of introgression and many hybrids between *Q. robur* and *Q. petraea* in many UK populations (Nocchi et al., 2021; Gathercole et al., 2026). Heavy population structure caused by the presence of multiple oak species will likely confound GWAS, therefore we performed a principal component analysis (PCA) with plink2 using the **reduced SNP set**, with the expectation that different oak species would be clearly separated in the PC space. To assist in our diagnosis, we used fastSTRUCTURE v1.0 (Raj et al., 2014) to infer admixture levels. Due to the high computational demands required to estimate admixture proportions, we randomly sampled 50,000 SNPs from the **reduced SNP set** which were used in fastSTRUCTURE with a simple prior, fivefold cross-validation, and number of ancestral populations (K) ranging from 1 to 10. We used fastSTUCTURE ChooseK script to select the best fitting K value. Clusters were ascribed to each species based on the geographical distribution of the “pure” samples i.e. those with less than 10% admixture (Lepais et al., 2009; Nocchi et al., 2021; Gathercole et al., 2026).

Next, we investigated population substructure in *Q. robur*, which was the most abundant oak species identified in the analyses above. After individuals with more than > 10% admixture were removed, we calculated fixation indices (F_ST_) between populations (*n* = 57), and pairwise kinship coefficients among individuals within populations using the **reduced SNP set**. For fixation indices we used the plink2 function –fst, and for kinship analysis we built a genetic relatedness matrix using plink2 --make-king function. Individuals and kinship coefficients were plotted using the Fruchterman-Reingold force-directed algorithm in the R package igraph v2.0.3 (Csárdi et al., 2025). Finally, linkage disequilibrium decay was calculated with PopLDdecay using the **gwas SNP set** (Zhang et al., 2019).

### SNP heritability

We estimated the SNP heritability 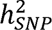 of phenotypic traits in LDAK v6 (Speed et al., 2017) using a restricted maximum likelihood (REML) regression. Phenotypes were regressed on the genomic relatedness matrix generated in the section above (GWAS, Equations 1 & 2) to estimate variance components, such that 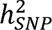, where 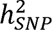 is the genetic variance and the residual, or environmental variance. To minimise potentially inflated estimates due to epistatic genetic effects and the shared environment in close relatives (Yang et al., 2017), pairs of individuals with kinship coefficients > 0.0884 (i.e. second-degree relations and above) were filtered out using plink2 --king-cutoff function. The function attempts to maximise the number of individuals remaining in the final set after pruning. Individuals from control sites were excluded from heritability models evaluating AOD and its separate symptoms (i.e. crown absolute transparency, stem bleeds, and presence of *Agrilus* exit holes). For mildew, only sites where at least one individual was found to be symptomatic were included (Supplementary Table 1). Final sample sizes for each phenotype are provided in Table 1.

### Genome-wide association study (GWAS)

We tested for associations between genetic markers and phenotypes in *Q. robur* using a linear mixed model (LMM) implemented in LDAK v6 (Speed et al., 2017). LMMs control for subtle population substructure and cryptic relatedness by fitting a genetic relatedness matrix (GRM) as a random effect in the model. While logistic regressions are more appropriate for binary traits, linear mixed models are a close approximation, particularly when the data lacks strong population stratification (Lee et al., 2011; Pirinen et al., 2013). Linear mixed models can be described by the formula:

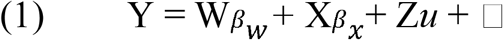

where Y is a vector of phenotypes, W is a matrix of covariates and β_w_ the vector of the unknown fixed effects, X is a matrix of genetic markers and β_x_ are its additive genetic effects, *u* are the random effects which follow a multivariate normal distribution (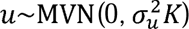), where *K* is the GRM and is a vector of the residual errors, which are also assumed to have a normal distribution (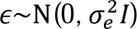) , where I is the identity matrix. The GRM generated in LDAK v6 has additional assumptions that take into account the effects of LD and MAF on SNP heritability (Speed et al., 2012, 2017). For example, estimated heritability can be inflated due to multi-tagging in LD regions, so LDAK sets 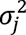 to 0 if SNP j has high correlation (r^2^ > 0.98) with SNP j’ and 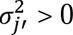. Moreover, because SNPs with low MAFs are expected to contribute less to heritability, LDAK weights SNPs as a function of MAF such that:

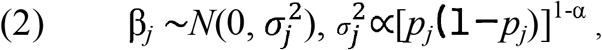

where β*_j_* is the effect of SNP_j_ and 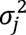 its variance, p is the MAF and α is the power scaling parameter. Many GWAS software developed for humans, such as GTCA (Yang et al., 2011), assume a power of -1 when constructing GRMs, but recently a power of -0.25 has been shown to be a good heritability approximation for many human traits (Speed et al., 2017). Most crop and animal research on the other hand, tend to assume 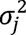 is directly proportional to MAF and therefore use a power of 0 (VanRaden, 2008). Preliminary analyses suggested α = -1 was best suited to our data and this value was therefore used throughout.

LMMs were built with the **gwas SNP set** as predictors and four phenotypes as the response, namely 1) the individual’s AOD diagnosis (0=control, 1=case), 2) presence of *Agrilus* exit holes (0=control, 1=case), 3) number of stem bleeds (discrete, with both active and inactive bleeds included), 4) absolute crown transparency, and 5) presence of mildew infection (0=control, 1=case). Absolute crown transparency was squared-root transformed to achieve normality, and number of stem bleeds was cube-root transformed to minimise skewness due to the excess of zeros and extreme values. Individuals from sites where we do not have historical records for presence of AOD (i.e. control sites), were excluded from the analyses of AOD and its symptoms. The remaining sites had a minimum of 4 individuals and a maximum of 306 (Supplementary Table 1).

Because AOD is a complex syndrome which has shown seasonal variation, we re-ran the analysis but this time including only individuals from sites we had historical data for the past 14 years (i.e. FR monitoring plots, *n* = 5). Individuals which were symptomatic for more than half of the monitoring duration (i.e. >= 8 years), or which died as a consequence of AOD, were considered cases (*n* = 95), and individuals who never manifested AOD symptoms were considered controls (*n* = 219). Individuals which manifested symptoms for less than 8 years (n=220) were not included. We reasoned that this approach would be able to detect extreme cases of susceptibility and resistance, which is not possible with samples for which we have only a single observation. While the statistical power is greatly reduced due to the much lower sample size (*n* = 314), we hypothesized that (1) the use of individuals with more extreme symptoms, and (2) the reduction in environmental noise due to the inclusion of fewer sites, could partially compensate for the lack of power.

All GWAS models included as covariates 6 environmental variables from the source site (Supplementary Table 2) previously shown to be associated with AOD (Brown et al., 2018, Shaw et al., 2025), the first 4 principal components summarising variation in the SNP data (Supplementary Figure 4), and a factor representing the different project tranches (n=4). Environmental variables were included to control for site specific conditions, PCs were included to correct for population substructure caused by the Blickling site, which we show to be driven by more than cryptic relatedness which LMMs can handle well (see Results), and tranche identity was included to control for variation in sampling year and potential biases due to differences in coverage, library preparation, and sequencing platform among tranches. For significance thresholds in the GWAS, we used three different approaches: (i) Bonferroni correction at a family-wise error rate of α = 0.05; (ii) a modified Bonferroni correction using the number of haplotype blocks rather than the number of SNPs (8,474,685); and (iii) the FDR controlled threshold at α = 0.05. Method (ii) seeks to correct for over-stringency in the regular Bonferroni correction taking into account that SNPs are not entirely independent due to linkage disequilibrium. Haplotype blocks were calculated with plink v1.9-170906 (Purcell et al., 2007) within a max distance of 1000kb. Manhattan plots and QQplots were generated using the R package topr v2.0.2 (Juliusdottir, 2023).

We searched for putative genes within 5kb intervals of SNPs most strongly associated with phenotypes in the GWAS using the *Q. robur* gene annotation file (Plomion et al., 2018), and a custom search function in the R v.4.2.2 environment (R Core Team, 2022). The function uses the gene position information in the annotation file to search for genes within 5kb upstream and downstream of the target SNP. If the function fails to assign any gene to the SNP within that distance, it repeats the search in incremental 5kb intervals until a gene is found. The function will then move to the next SNP until all selected SNPs are ascribed a putative gene. If more than one gene is found within the 5kb interval, both are reported in the output file.

Finally, we investigated whether SNPs and nearby genes associated with AOD or mildew were located within regions of particularly low differentiation between *Q. robur* and *Q. petraea*. Introgression between the two species in our populations is very common (Gathercole et al., 2026), so we speculated alleles from *Q. petraea*, which is more tolerant to mildew infection (Marçais & Desprez-Loustau, 2014; Lonsdale, 2016) and also to dry conditions (Eaton et al., 2016), could help *Q. robur* cope with fungal load and novel environmental risk factors associated with AOD e.g. recent increases in temperature in the southern regions of the UK. We checked whether genes identified here as associated with mildew or AOD were included in the list of genes located in regions of the genome with high differentiation between *Q. robur* and *Q. petraea* (Gathercole et al., 2026). Additionally, we investigated whether the alleles found to be associated with mildew or AOD differed in frequency between populations of *Q. petraea* and *Q. robur* using the plink2 function –freq.

### Genomic prediction

To test whether we could accurately predict phenotypic traits using genetic markers, we built elastic net (ENET) prediction models using LDAK v6 (Hof & Speed, 2025). ENET is a penalized regression which combines the ridge and LASSO penalties (Zou & Hastie, 2005) and can therefore capture both sparse (few large-effect SNPs) and polygenic (many small-effect SNPs) architectures, offering improved prediction accuracy, especially when the underlying genetic architecture is unknown. In the context of genomic prediction, the formula can be expressed as follows:

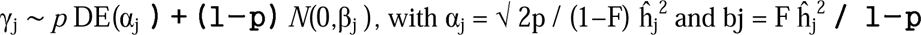

where DE(α_j_) stands for a double exponential distribution with rate α_j._ The relative contributions of the double exponential and normal distributions are regulated by the p and F parameters, respectively. LDAK chooses suitable parameters by a cross-validation procedure using a subset of samples, and chooses the model which minimizes the mean-squared error. If the cross-validation step is omitted, LDAK generates 10 models with differing parameters, each trained with 100% of samples.

Here we use a classical 10-fold cross-validation procedure (Daetwyler et al., 2013; Zhou et al., 2017) involving randomly distributing samples into a training set containing 90% of samples and a test set with the remaining 10%. The procedure is repeated 10x until all samples are included in both sets. We fitted all ∼8 million SNPs in each model, since ENET models have a shrinking parameter. We did not do this procedure for mildew as there were not enough cases for the allocation of a sufficient number of affected individuals to test sets.

For AOD occurrence, we investigated whether models trained with markers discovered in a GWAS could lead to more accurate predictions despite the lower density (e.g. Hiraoka et al., 2018; Stocks et al., 2019). We performed a GWAS as in the section above, but this time excluding one of the randomised sets from the model which had led to particularly high accuracy. The most significant predictors (i.e. those with the lowest p-values) discovered in the GWAS were fitted incrementally (i.e. 100, 500, 1k, 5k, 10k, 50k, 100k, 500k) in different GP models. Finally, to confirm whether models trained with GWAS markers performed better than randomly selected markers, we repeated the incremental strategy but this time selecting markers randomly with the plink2 function --thin-count. The accuracy of model predictions was determined by Pearson’s correlation between the genomic estimated breeding values (GEBVs) and the observed phenotypes of the test data. The same training and test sets were used in all models so that they were comparable.

## DATA ACCESSIBILITY

Raw reads generated in this project were uploaded to the NCBI public database (BioProject reference: PRJNA816149).

## ACKNOWLEDGEMENTS

This project was funded by the Department for Environment, Food & Rural Affairs via the Center for Forest Protection (Ref. CFP2211). This research utilised Queen Mary’s Apocrita HPC facility, supported by QMUL Research-IT (http://doi.org/10.5281/zenodo.438045). We thank landowners for access to their trees. We thank the FR technical services unit for sample collection and phenotyping.

## AUTHOR INFORMATION

### Contributions

R.C. conducted analyses and DNA lab work and wrote the paper. L.G. and G. N. contributed data and commented on the paper. F. C-D. conducted lab work. N.B. contributed site and phenotype data. S.D. and R. B. conceived the study, gained funding, led the project and wrote the paper.

### Corresponding authors

Correspondence to Richard Buggs and Rômulo Carleial

## REFERENCES

Andrews, S. (2010). FastQC: A Quality Control Tool for High Throughput Sequence Data [Online]

Ayres, P. G. (1976). Natural resistance to oak mildew. Arboricultural Journal, 3(1), 23–29. 10.1080/03071375.1976.10590474

Barrès, B., Dutech, C., Saint-Jean, G., Bodénès, C., Burban, C., Fiévet, V., … & Desprez-Loustau, M. L. (2024). Demographic and genetic impacts of powdery mildew in a young oak (*Quercus robur* L.) cohort. Annals of Forest Science, 81(1), 44. 10.1186/s13595-024-01259-2

Bartholomé, J., Brachi, B., Marçais, B., Mougou Hamdane, A., Bodénès, C., Plomion, C., … & Desprez Loustau, M. L. (2020). The genetics of exapted resistance to two exotic pathogens in pedunculate oak. New Phytologist, 226(4), 1088–1103. 10.1111/nph.16319

Beatty, G. E., Montgomery, W. I., Spaans, F., Tosh, D. G., & Provan, J. (2016). Pure species in a continuum of genetic and morphological variation: sympatric oaks at the edge of their range. Annals of Botany, 117(4), 541–549. 10.1093/aob/mcw002

Beaulieu, J., Doerksen, T., Clément, S., MacKay, J., & Bousquet, J. (2014). Accuracy of genomic selection models in a large population of open-pollinated families in white spruce. Heredity, 113(4), 343–352. 10.1038/hdy.2014.36

Brady, C., Arnold, D., McDonald, J., & Denman, S. (2017). Taxonomy and identification of bacteria associated with acute oak decline. World Journal of Microbiology and Biotechnology, 33, 1–11. 10.1007/s11274-017-2296-4

Brown, N., Inward, D. J., Jeger, M., & Denman, S. (2015). A review of Agrilus biguttatus in UK forests and its relationship with acute oak decline. Forestry: An International Journal of Forest Research, 88(1), 53–63. 10.1093/forestry/cpu039

Brown, N., Jeger, M., Kirk, S., Xu, X., & Denman, S. (2016). Spatial and temporal patterns in symptom expression within eight woodlands affected by Acute Oak Decline. Forest Ecology and Management, 360, 97–109. 10.1016/j.foreco.2015.10.026

Brown, N., Jeger, M., Kirk, S., Williams, D., Xu, X., Pautasso, M., & Denman, S. (2017). Acute oak decline and Agrilus biguttatus: The co-occurrence of stem bleeding and D-shaped emergence holes in Great Britain. Forests, 8(3), 87. 10.3390/f8030087

Brown, N., Vanguelova, E., Parnell, S., Broadmeadow, S., & Denman, S. (2018). Predisposition of forests to biotic disturbance: Predicting the distribution of Acute Oak Decline using environmental factors. Forest Ecology and Management, 407, 145–154. 10.1016/j.foreco.2017.10.054

Carleial, R., Charters, M. D., Finzgar, D., Swift, P., Anthoney, E., Sahlstedt, E., … & Buggs, R. (2025). Genetic basis of traits and local adaptation in UK silver birch. bioRxiv, 2025–07. 10.1101/2025.07.04.662749

Carluccio, G., Benigno, A., Panzavolta, T., Vergine, M., De Bellis, L., Luvisi, A., & Moricca, S. (2025). Understanding Oak Decline in Europe: Ecological Factors, Symptoms, Causative Agents, and Management Strategies. Plant Disease, 109(9), 1805–1823. 10.1094/PDIS-11-24-2401-FE

Chai, L. X., Dong, K., Liu, S. Y., Zhang, Z., Zhang, X. P., Tong, X., … & Wang, X. B. (2020). A putative nuclear copper chaperone promotes plant immunity in Arabidopsis. Journal of Experimental Botany, 71(20), 6684–6696. 10.1093/jxb/eraa401

Chang, C. C., Chow, C. C., Tellier, L. C., Vattikuti, S., Purcell, S. M., & Lee, J. J. (2015). Second-generation PLINK: rising to the challenge of larger and richer datasets. Gigascience, 4(1), s13742–015. 10.1186/s13742-015-0047-8

Crampton, B. G., Plummer, S. J., Kaczmarek, M., McDonald, J. E., & Denman, S. (2020). A multiplex real time PCR assay enables simultaneous rapid detection and quantification of bacteria associated with acute oak decline. Plant Pathology, 69(7), 1301–1310. 10.1111/ppa.13203

Crossa, J., Pérez-Rodríguez, P., Cuevas, J., Montesinos-López, O., Jarquín, D., De Los Campos, G., … & Varshney, R. K. (2017). Genomic selection in plant breeding: methods, models, and perspectives. Trends in plant science, 22(11), 961–975. 10.1016/j.tplants.2017.08.011

Csárdi G, Nepusz T, Traag V, Horvát Sz, Zanini F, Noom D, Müller K (2025). igraph: Network Analysis and Visualization in R_. doi:10.5281/zenodo.7682609 <10.5281/zenodo.7682609>, R package version 2.0.3, <https://CRAN.R-project.org/package=igraph>.

Daetwyler, H. D., Calus, M. P., Pong-Wong, R., de Los Campos, G., & Hickey, J. M. (2013). Genomic prediction in animals and plants: simulation of data, validation, reporting, and benchmarking. Genetics, 193(2), 347–365. 10.1534/genetics.112.147983

Danecek, P., Auton, A., Abecasis, G., Albers, C. A., Banks, E., DePristo, M. A., … & 1000 Genomes Project Analysis Group. (2011). The variant call format and VCFtools. Bioinformatics, 27(15), 2156–2158. 10.1093/bioinformatics/btr330

Demeter, L., Molnár, Á. P., Öllerer, K., Csóka, G., Kiš, A., Vadász, C., … & Molnár, Z. (2021). Rethinking the natural regeneration failure of pedunculate oak: The pathogen mildew hypothesis. Biological Conservation, 253, 108928. 10.1016/j.biocon.2020.108928

Denman, S., Webber, J., 2009. Oak declines: new definitions and new episodes in Britain. Quart. J. Forestry 103, 285– 290.

Denman, S., Brown, N., Kirk, S., Jeger, M., & Webber, J. (2014). A description of the symptoms of Acute Oak Decline in Britain and a comparative review on causes of similar disorders on oak in Europe. Forestry: An International Journal of Forest Research, 87(4), 535–551. 10.1093/forestry/cpu010

Denman, S., Doonan, J., Ransom-Jones, E., Broberg, M., Plummer, S., Kirk, S., … & McDonald, J. E. (2018). Microbiome and infectivity studies reveal complex polyspecies tree disease in Acute Oak Decline. The ISME journal, 12(2), 386–399. 10.1038/ismej.2017.170

Denman, S., Brown, N., Vanguelova, E., & Crampton, B. (2022). Temperate Oak Declines: Biotic and abiotic predisposition drivers. Forest microbiology, 239–263. 10.1016/B978-0-323-85042-1.00020-3

DePristo, M. A., Banks, E., Poplin, R., Garimella, K. V., Maguire, J. R., Hartl, C., … & Daly, M. J. (2011). A framework for variation discovery and genotyping using next-generation DNA sequencing data. Nature genetics, 43(5), 491–498. 10.1038/ng.806

Desprez-Loustau, M. L., Massot, M., Feau, N., Fort, T., de Vicente, A., Torés, J. A., & Ortuño, D. F. (2017). Further support of conspecificity of oak and mango powdery mildew and first report of *Erysiphe quercicola* and *Erysiphe alphitoides* on mango in mainland Europe. Plant Disease, 101(7), 1086–1093. 10.1094/PDIS-01-17-0116-RE

Doonan, J., Denman, S., Pachebat, J. A., & McDonald, J. E. (2019). Genomic analysis of bacteria in the Acute Oak Decline pathobiome. Microbial genomics, 5(1), e000240. 10.1099/mgen.0.000240

Doonan, J. M., Broberg, M., Denman, S., & McDonald, J. E. (2020). Host–microbiota–insect interactions drive emergent virulence in a complex tree disease. Proceedings of the Royal Society B, 287(1933), 20200956. 10.1098/rspb.2020.0956

Eaton, E. G. S. D. J., Caudullo, G., Oliveira, S., & De Rigo, D. (2016). *Quercus robur* and *Quercus petraea* in Europe: distribution, habitat, usage and threats. European atlas of forest tree species, 14, 160–163

Edelman, N. B., & Mallet, J. (2021). Prevalence and adaptive impact of introgression. Annual review of genetics, 55(1), 265–283. 10.1146/annurev-genet-021821-020805

Finch, J. P., Brown, N., Beckmann, M., Denman, S., & Draper, J. (2021). Index measures for oak decline severity using phenotypic descriptors. Forest Ecology and Management, 485, 118948. 10.1016/j.foreco.2021.118948

Fletcher, K., Han, R., Smilde, D., & Michelmore, R. (2022). Variance of allele balance calculated from low coverage sequencing data infers departure from a diploid state. BMC bioinformatics, 23(1), 150. 10.1186/s12859-022-04685-z

Forestry Commission (2023). Acute Oak Decline (AOD): Incidence and Distribution. https://www.forestresearch.gov.uk/tools-and-resources/fthr/pest-and-disease-resources/acute-oak-decline/acute-oak-decline-aod-incidence-and-distribution/#:~:text=AOD%20has%20been%20found%20mainly,occurrences%20along%20the%20Welsh%20borders. Accessed 10/02/2025.

Gallagher, M. D., & Chen-Plotkin, A. S. (2018). The post-GWAS era: from association to function. The American Journal of Human Genetics, 102(5), 717–730. 10.1016/j.ajhg.2018.04.002

Gathercole, L. A., Nocchi, G., Brown, N., Coker, T. L., Plumb, W. J., Stocks, J. J., … & Buggs, R. J. (2021). Evidence for the widespread occurrence of bacteria implicated in acute oak decline from incidental genetic sampling. Forests, 12(12), 1683. 10.3390/f12121683

Gathercole, L. A., Carleial, R., Brown, N., Deman, S., Wu, E., Nichols, R. A., & Buggs, R. J. A. (2026). Genomic diversity of British native oaks: species differentiation, hybridisation and triploidy. BioRxiv 708962

Gienapp, P., Fior, S., Guillaume, F., Lasky, J. R., Sork, V. L., & Csilléry, K. (2017). Genomic quantitative genetics to study evolution in the wild. Trends in Ecology & Evolution, 32(12), 897–908. 10.1016/j.tree.2017.09.004

Gil-Pelegrín, E., Peguero-Pina, J. J., & Sancho-Knapik, D. (2017). Oaks and people: a long journey together. In: Gil-Pelegrín, E., Peguero-Pina, J., Sancho-Knapik, D. (eds) Oaks Physiological Ecology. Exploring the Functional Diversity of Genus Quercus L. Tree Physiology, vol 7. Springer, Cham. 10.1007/978-3-319-69099-5_1

Grattapaglia, D. (2017). Status and Perspectives of Genomic Selection in Forest Tree Breeding. In: Varshney, R., Roorkiwal, M., Sorrells, M. (eds) Genomic Selection for Crop Improvement. Springer, Cham. 10.1007/978-3-319-63170-7_9

Grattapaglia, D. (2022). Twelve years into genomic selection in forest trees: climbing the slope of enlightenment of marker assisted tree breeding. Forests, 13(10), 1554. 10.3390/f13101554

Guo, C., Luo, C., Guo, L., Li, M., Guo, X., Zhang, Y., … & Chen, L. (2016). OsSIDP366, a DUF1644 gene, positively regulates responses to drought and salt stresses in rice. Journal of integrative plant biology, 58(5), 492–502. 10.1111/jipb.12376

Hill, W. G. (2014). Applications of population genetics to animal breeding, from Wright, Fisher and Lush to genomic prediction. Genetics, 196(1), 1–16. 10.1534/genetics.112.147850

Hiraoka, Y., Fukatsu, E., Mishima, K., Hirao, T., Teshima, K. M., Tamura, M., … & Watanabe, A. (2018). Potential of genome-wide studies in unrelated plus trees of a coniferous species, *Cryptomeria japonica* (Japanese cedar). Frontiers in Plant Science, 9, 1322. 10.3389/fpls.2018.01322

Hof, J. P., & Speed, D. (2025). LDAK-KVIK performs fast and powerful mixed-model association analysis of quantitative and binary phenotypes. Nature Genetics, 57(9), 2116-2123. 10.1038/s41588-025-02286-z5

Isik, F., Cooperative Tree Improvement Program, Shalizi, M. N., & Walker, T. D. (2026). Genomic selection validated across two generations of loblolly pine breeding. bioRxiv, 2026–01. 10.64898/2026.01.22.701135

Jensen, J., Larsen, A., Nielsen, L. R., & Cottrell, J. (2009). Hybridization between Quercus robur and Q. petraea in a mixed oak stand in Denmark. Annals of Forest Science, 66(7), 1–12. 10.1051/forest/2009058

Juliusdottir, T. (2023). topr: an R package for viewing and annotating genetic association results. BMC bioinformatics, 24(1), 268. 10.1186/s12859-023-05301-4

Knaus, B. J., & Grünwald, N. J. (2017). vcfr: a package to manipulate and visualize variant call format data in R. Molecular ecology resources, 17(1), 44–53. 10.1111/1755-0998.12549

Kremer, A., & Hipp, A. L. (2020). Oaks: an evolutionary success story. New Phytologist, 226(4), 987–1011. 10.1111/nph.16274

Lang T, Abadie P, Léger V, Decourcelle T, Frigerio J-M, Burban C et al (2021) High-quality SNPs from genic regions highlight introgression patterns among European white oaks (*Quercus petraea* and *Q. robur)*. https://www.biorxiv.org/content/10.1101/388447v4.full.pdf

Lebedev, V. G., Lebedeva, T. N., Chernodubov, A. I., & Shestibratov, K. A. (2020). Genomic selection for forest tree improvement: Methods, achievements and perspectives. Forests, 11(11), 1190. 10.3390/f11111190

Lee, S. H., Wray, N. R., Goddard, M. E., & Visscher, P. M. (2011). Estimating missing heritability for disease from genome-wide association studies. The American Journal of Human Genetics, 88(3), 294–305. https://10.1016/j.ajhg.2011.02.002

Lepais, O., Petit, R. J., Guichoux, E., Lavabre, J. E., Alberto, F., Kremer, A., & Gerber, S. (2009). Species relative abundance and direction of introgression in oaks. Molecular Ecology, 18(10), 2228–2242. 10.1111/j.1365-294X.2009.04137.x

Leroy, T., Louvet, J. M., Lalanne, C., Le Provost, G., Labadie, K., Aury, J. M., … & Kremer, A. (2020). Adaptive introgression as a driver of local adaptation to climate in European white oaks. New Phytologist, 226(4), 1171–1182. 10.1111/nph.16095

Li, H., & Durbin, R. (2009). Fast and accurate short read alignment with Burrows–Wheeler transform. Bioinformatics, 25(14), 1754–1760. 10.1093/bioinformatics/btp324

Li, H., Handsaker, B., Wysoker, A., Fennell, T., Ruan, J., Homer, N., … & 1000 Genome Project Data Processing Subgroup. (2009). The sequence alignment/map format and SAMtools. bioinformatics, 25(16), 2078–2079. 10.1093/bioinformatics/btp352

Li, T., Liao, K., Xu, X., Gao, Y., Wang, Z., Zhu, X., … & Xuan, Y. (2017). Wheat ammonium transporter (AMT) gene family: diversity and possible role in host–pathogen interaction with stem rust. Frontiers in plant science, 8, 1637.

Li, C. Y., Zhang, N., Bin, G. U. A. N., Zhou, Z. Q., & Mei, F. Z. (2019). Reactive oxygen species are involved in cell death in wheat roots against powdery mildew. Journal of Integrative Agriculture, 18(9), 1961–1970. 10.1016/S2095-3119(18)62092-1

Lonsdale, D. (2015). Review of oak mildew, with particular reference to mature and veteran trees in Britain. Arboricultural Journal, 37(2), 61–84. 10.1080/03071375.2015.1039839

Lonsdale, D. (2016). Powdery mildew of oak: a familiar sight with some hidden surprises. The ARB magazine, 43, 48–52.

Marçais, B., & Desprez-Loustau, M. L. (2014). European oak powdery mildew: impact on trees, effects of environmental factors, and potential effects of climate change. Annals of Forest Science, 71, 633–642. 10.1007/s13595-012-0252-x

Marees, A. T., De Kluiver, H., Stringer, S., Vorspan, F., Curis, E., Marie Claire, C., & Derks, E. M. (2018). A tutorial on conducting genome wide association studies: Quality control and statistical analysis. International journal of methods in psychiatric research, 27(2), e1608. 10.1002/mpr.1608

McGaugh, S. E., Lorenz, A. J., & Flagel, L. E. (2021). The utility of genomic prediction models in evolutionary genetics. Proceedings of the Royal Society B, 288(1956), 20210693. 10.1098/rspb.2021.0693

McKenna, A., Hanna, M., Banks, E., Sivachenko, A., Cibulskis, K., Kernytsky, A., … & DePristo, M. A. (2010). The Genome Analysis Toolkit: a MapReduce framework for analyzing next-generation DNA sequencing data. Genome research, 20(9), 1297–1303. 10.1101/gr.107524.110

Meuwissen, T. H., Hayes, B. J., & Goddard, M. (2001). Prediction of total genetic value using genome-wide dense marker maps. genetics, 157(4), 1819–1829. 10.1093/genetics/157.4.1819

Mitchell, R. J., Bellamy, P. E., Ellis, C. J., Hewison, R. L., Hodgetts, N. G., Iason, G. R., … & Taylor, A. F. S. (2019). OakEcol: A database of Oak-associated biodiversity within the UK. Data in brief, 25, 104120. 10.5285/22b3d41e-7c35-4c51-9e55-0f47bb845202

Müller, B. S., Neves, L. G., de Almeida Filho, J. E., Resende, M. F., Muñoz, P. R., Dos Santos, P. E., … & Grattapaglia, D. (2017). Genomic prediction in contrast to a genome-wide association study in explaining heritable variation of complex growth traits in breeding populations of Eucalyptus. BMC genomics, 18, 1–17. 10.1186/s12864-017-3920-2

Nocchi, G., Brown, N., Coker, T. L., Plumb, W. J., Stocks, J. J., Denman, S., & Buggs, R. J. (2021). Genomic structure and diversity of oak populations in British parklands. *Plants, People*, Planet, 4(2), 167–181. 10.1002/ppp3.10229

Padmanabhan, M., Cournoyer, P., & Dinesh Kumar, S. P. (2009). The leucine rich repeat domain in plant innate immunity: a wealth of possibilities. Cellular microbiology, 11(2), 191–198. 10.1111/j.1462-5822.2008.01260.x

Petit, R. J., Bodénès, C., Ducousso, A., Roussel, G., & Kremer, A. (2004). Hybridization as a mechanism of invasion in oaks. New Phytologist, 161(1), 151–164. 10.1046/j.1469-8137.2003.00944.x

Pino Del Carpio, D., Lozano, R., Wolfe, M.D., Jannink, JL. (2019). Genome-Wide Association Studies and Heritability Estimation in the Functional Genomics Era. In: Rajora, O. (eds) Population Genomics. Concepts, Approaches and Applications. Springer, Cham. 10.1007/13836_2018_12

Pirinen, M., Donnelly, P., & Spencer, C. C. (2013). Efficient computation with a linear mixed model on large-scale data sets with applications to genetic studies. The Annals of Applied Statistics, 369–390. 10.1214/12-AOAS586

Plomion, C., Aury, J. M., Amselem, J., Leroy, T., Murat, F., Duplessis, S., … & Salse, J. (2018). Oak genome reveals facets of long lifespan. Nature plants, 4(7), 440–452. 10.1038/s41477-018-0172-3

Plumb, W. J., Coker, T. L., Stocks, J. J., Woodcock, P., Quine, C. P., Nemesio Gorriz, M., … & Buggs, R. J. (2020). The viability of a breeding programme for ash in the British Isles in the face of ash dieback. Plants, People, Planet, 2(1), 29–40. 10.1002/ppp3.10060

Poland, J., & Rutkoski, J. (2016). Advances and challenges in genomic selection for disease resistance. Annual review of phytopathology, 54(1), 79–98. 10.1146/annurev-phyto-080615-100056

Purcell, S., Neale, B., Todd-Brown, K., Thomas, L., Ferreira, M. A., Bender, D., … & Sham, P. C. (2007). PLINK: a tool set for whole-genome association and population-based linkage analyses. The American journal of human genetics, 81(3), 559–575. 10.1086/519795

R Core Team (2022). R: A language and environment for statistical computing. R Foundation for Statistical Computing, Vienna, Austria. URL https://www.R-project.org/.

Raj, A., Stephens, M., & Pritchard, J. K. (2014). fastSTRUCTURE: variational inference of population structure in large SNP data sets. Genetics, 197(2), 573–589. 10.1534/genetics.114.164350

Reed, E., Nunez, S., Kulp, D., Qian, J., Reilly, M. P., & Foulkes, A. S. (2015). A guide to genome wide association analysis and post analytic interrogation. Statistics in medicine, 34(28), 3769–3792. 10.1002/sim.6605

Reed, K., Denman, S., Leather, S. R., Forster, J., & Inward, D. J. (2018). The lifecycle of *Agrilus biguttatus*: the role of temperature in its development and distribution, and implications for Acute Oak Decline. Agricultural and Forest Entomology, 20(3), 334–346. 10.1111/afe.12266

Rey, P., & Tarrago, L. (2018). Physiological roles of plant methionine sulfoxide reductases in redox homeostasis and signaling. Antioxidants, 7(9), 114. 10.3390/antiox7090114

Samorè, A. B., & Fontanesi, L. (2016). Genomic selection in pigs: state of the art and perspectives. Italian Journal of Animal Science, 15(2), 211–232. 10.1080/1828051X.2016.1172034

Sanchez Lucas, R., Bosanquet, J. L., Henderson, J., Catoni, M., Pastor, V., & Luna, E. (2025). Elicitor specific mechanisms of defence priming in oak seedlings against powdery mildew. Plant, Cell & Environment, 48(6), 4455–4474. 10.1111/pce.15419

Scarlett, K., Denman, S., Clark, D. R., Forster, J., Vanguelova, E., Brown, N., & Whitby, C. (2021). Relationships between nitrogen cycling microbial community abundance and composition reveal the indirect effect of soil pH on oak decline. The ISME Journal, 15(3), 623–635. 10.1038/s41396-020-00801-0

Shaw, L. J., Rabiey, M., Garcia, M. S., Clarke, T., Broome, A., Corbett, L. J., … & Ray, D. (2025). The cause-effect conundrum of local-scale site and soil factors in acute oak decline (AOD). bioRxiv, 2025-01. 10.1101/2025.01.24.634765

Shen, C., Yuan, J., Ou, X., Ren, X., & Li, X. (2021). Genome-wide identification of alcohol dehydrogenase (ADH) gene family under waterlogging stress in wheat (Triticum aestivum). PeerJ, 9, e11861. 10.7717/peerj.11861

Speed, D., Hemani, G., Johnson, M. R., & Balding, D. J. (2012). Improved heritability estimation from genome-wide SNPs. The American Journal of Human Genetics, 91(6), 1011–1021. 10.1016/j.ajhg.2012.10.010

Speed, D., Cai, N., Ucleb Consortium, Johnson, M. R., Nejentsev, S., & Balding, D. J. (2017). Reevaluation of SNP heritability in complex human traits. Nature genetics, 49(7), 986–992. 10.1038/ng.3865 https://doi.org/10.1016/j.ajhg.2012.10.010

Stocks, J. J., Metheringham, C. L., Plumb, W. J., Lee, S. J., Kelly, L. J., Nichols, R. A., & Buggs, R. J. (2019). Genomic basis of European ash tree resistance to ash dieback fungus. Nature ecology & evolution, 3(12), 1686–1696. 10.1038/s41559-019-1036-6

Tade, B., & Melesse, A. (2024). A review on the application of genomic selection in the improvement of dairy cattle productivity. Ecological Genetics and Genomics, 31, 100257. 10.1016/j.egg.2024.100257

Thomas, F. M., Blank, R., & Hartmann, G. (2002). Abiotic and biotic factors and their interactions as causes of oak decline in Central Europe. Forest Pathology, *32*(4 5), 277-307. 10.1046/j.1439-0329.2002.00291.x

Thomas, F. M. (2008). Recent advances in cause-effect research on oak decline in Europe. CABI Reviews, (2008), 12- pp. 10.1079/PAVSNNR20083037

Tibbs Cortes, L., Zhang, Z., & Yu, J. (2021). Status and prospects of genome wide association studies in plants. The plant genome, 14(1), e20077. 10.1002/tpg2.20077

Unno, H., Uchida, T., Sugawara, H., Kurisu, G., Sugiyama, T., Yamaya, T., … & Kusunoki, M. (2006). Atomic structure of plant glutamine synthetase: a key enzyme for plant productivity. Journal of Biological Chemistry, 281(39), 29287–29296. 10.1074/jbc.M601497200

VanRaden, P. M. (2008). Efficient methods to compute genomic predictions. Journal of dairy science, 91(11), 4414–4423. 10.3168/jds.2007-0980

Vargas, J. A., Leonardo, D. A., D’Muniz Pereira, H., Lopes, A. R., Rodriguez, H. N., Cobos, M., … & Garratt, R. C. (2022). Structural characterization of l-galactose dehydrogenase: An essential enzyme for vitamin c biosynthesis. Plant and Cell Physiology, 63(8), 1140–1155. 10.1093/pcp/pcac090

Yang, J., Lee, S. H., Goddard, M. E., & Visscher, P. M. (2011). GCTA: a tool for genome-wide complex trait analysis. The American Journal of Human Genetics, 88(1), 76–82. 10.1016/j.ajhg.2010.11.011

Yang, J., Zeng, J., Goddard, M. E., Wray, N. R., & Visscher, P. M. (2017). Concepts, estimation and interpretation of SNP-based heritability. Nature genetics, 49(9), 1304–1310. 10.1038/ng.3941

Yoo, J. Y., Ko, K. S., Vu, B. N., Lee, Y. E., Yoon, S. H., Pham, T. T., … & Lee, K. O. (2021). N-acetylglucosaminyltransferase II is involved in plant growth and development under stress conditions. Frontiers in Plant Science, 12, 761064. 10.3389/fpls.2021.761064

Zarnoch, S. J., Bechtold, W. A., & Stolte, K. W. (2004). Using crown condition variables as indicators of forest health. Canadian Journal of Forest Research, 34(5), 1057–1070. 10.1139/x03-277

Zhang, C., Dong, S. S., Xu, J. Y., He, W. M., & Yang, T. L. (2019). PopLDdecay: a fast and effective tool for linkage disequilibrium decay analysis based on variant call format files. Bioinformatics, 35(10), 1786–1788. 10.1093/bioinformatics/bty875

Zhou, Y., Vales, M. I., Wang, A., & Zhang, Z. (2017). Systematic bias of correlation coefficient may explain negative accuracy of genomic prediction. Briefings in bioinformatics, 18(5), 744–753. 10.1093/bib/bbw064

Zou, H., & Hastie, T. (2005). Regularization and variable selection via the elastic net. Journal of the Royal Statistical Society Series B: Statistical Methodology, 67(2), 301–320. 10.1111/j.1467-9868.2005.00527.x

