## Supplementary Material for "Genomic basis for resistance to acute oak decline and mildew infection in English oak"

**SUPPLEMENTARY FIGURES**

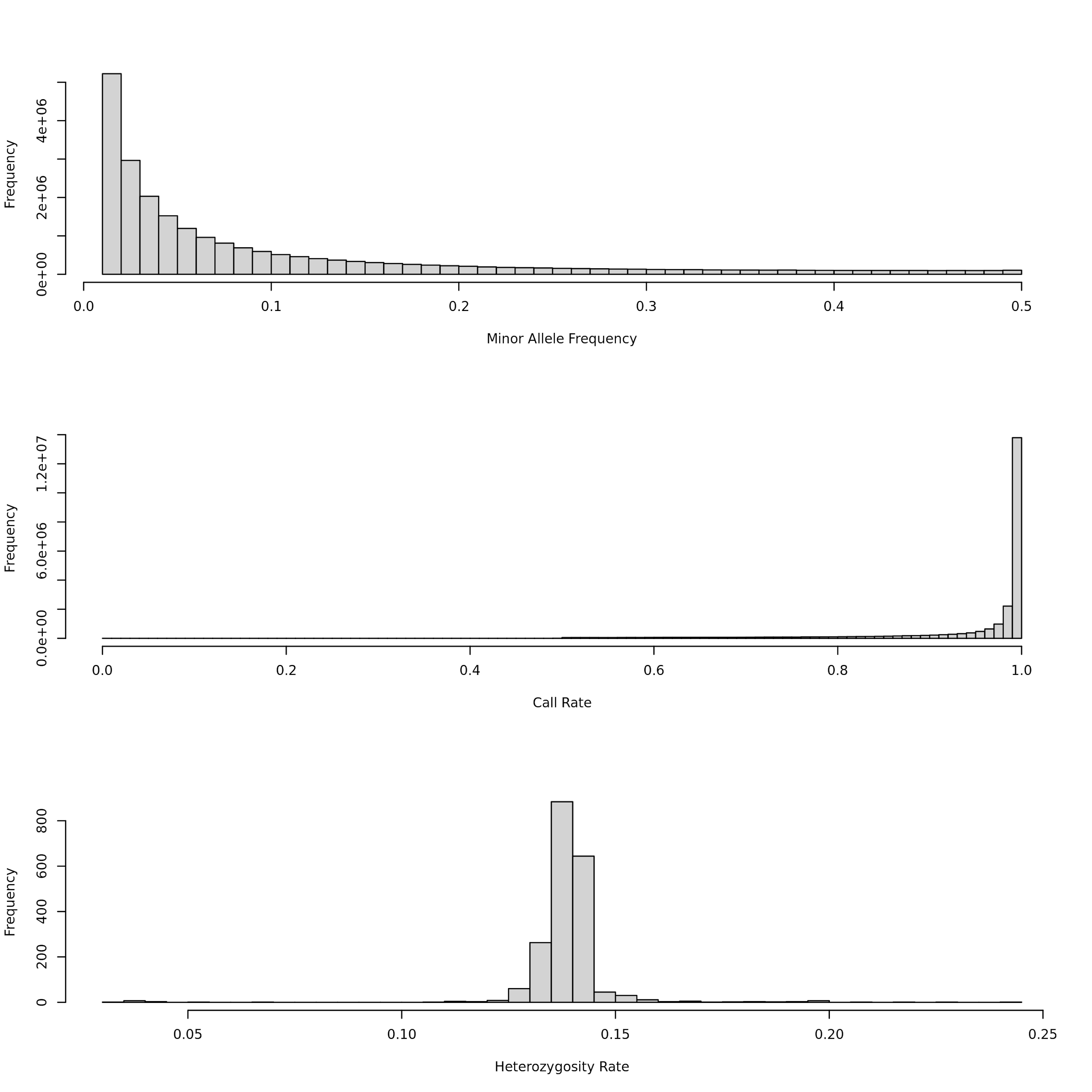
**Supplementary** **Fig. 1 - Summary statistics for SNP data including all 1996 *Quercus spp*. samples.** Statistics were calculated after three steps of hard-filtering (see Methods). SNPs were posteriorly filtered for minor allele frequencies (<0.05) and deviations from Hardy-Weinberg equilibrium (p < 1^-50^). Individuals strongly deviating (±2 SD) from the mean heterozygosity rate (bottom panel) were also removed.

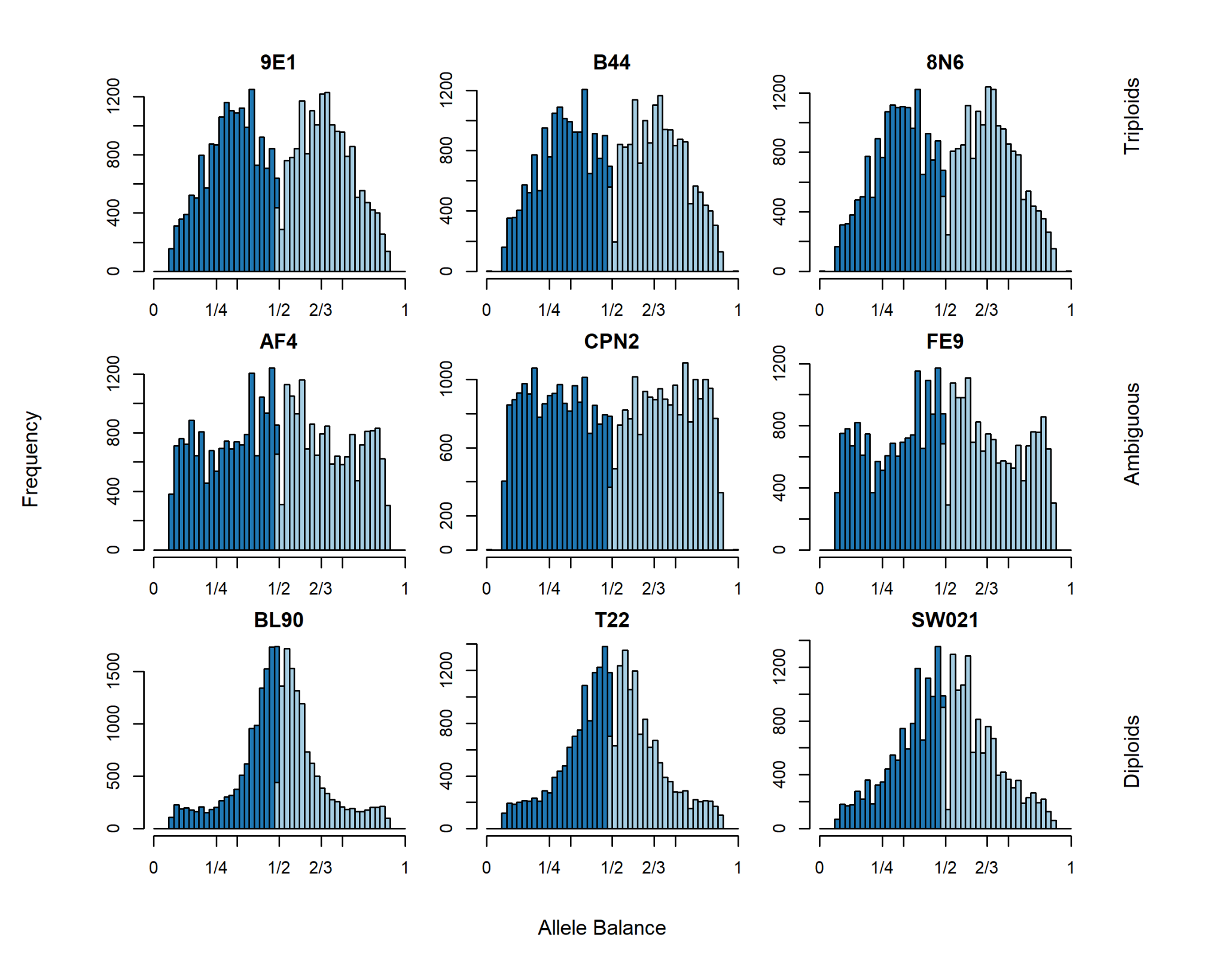

**Supplementary Figure 2 - Bar charts showing the allele balance proportions of six oak samples.** Samples on the top row have the typical allele distribution found in triploids. The middle row has individuals for which ploidy could not be determined. Bottom row samples have a typical diploid allele balance with the majority of loci having a roughly equal number of reads supporting each allele.

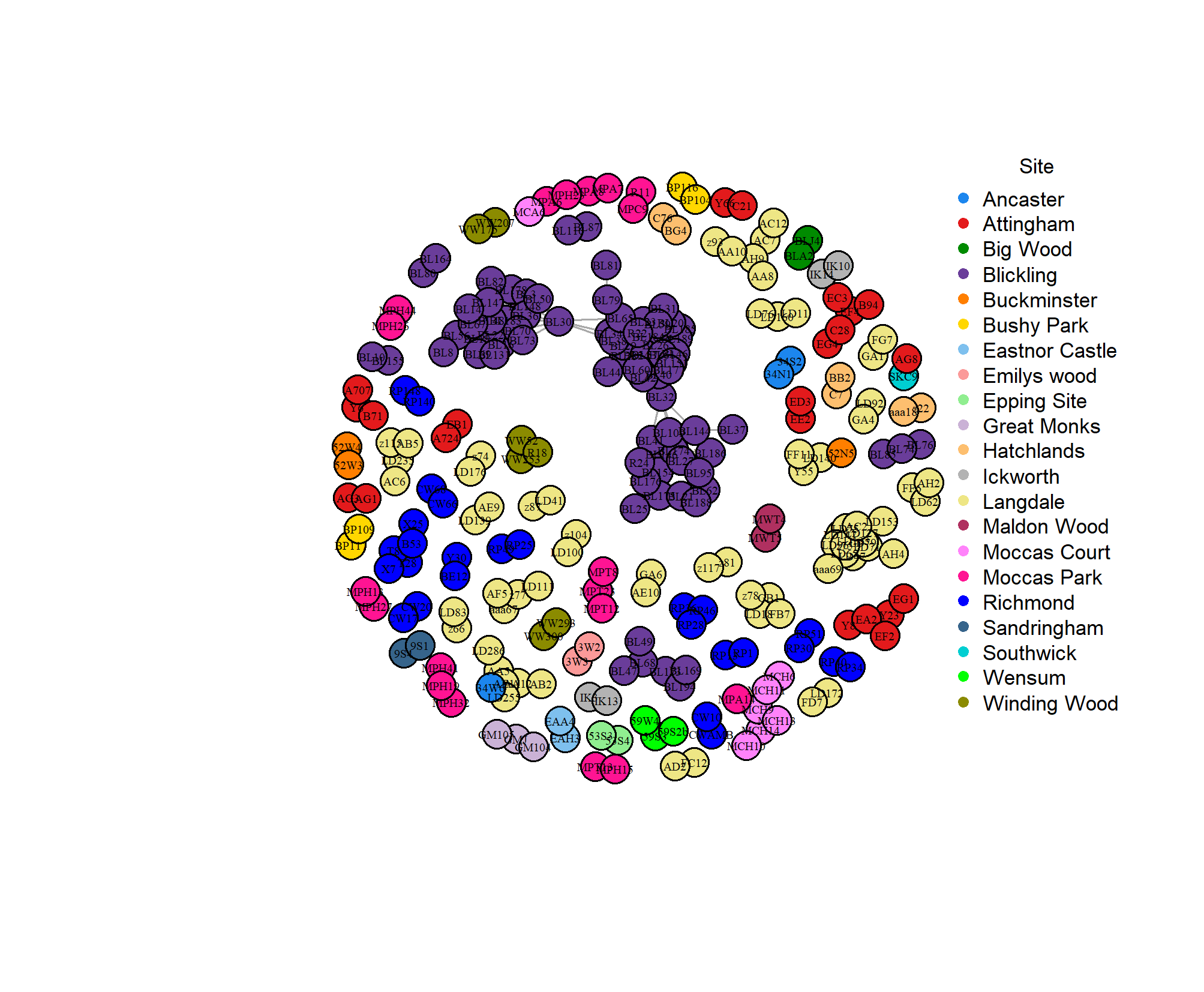

**Supplementary Fig 3 - Network visualization of kinship relations among *Quercus robur* individuals used in this study.** Nodes are colour coded by collection site and represent individuals, and edge weights correspond to pairwise kinship coefficients. Clusters indicate groups of closely related individuals (1^st^ and 2^nd^ degree). Blickling (purple) had a disproportionally large number of closely related individuals compared to other sites.

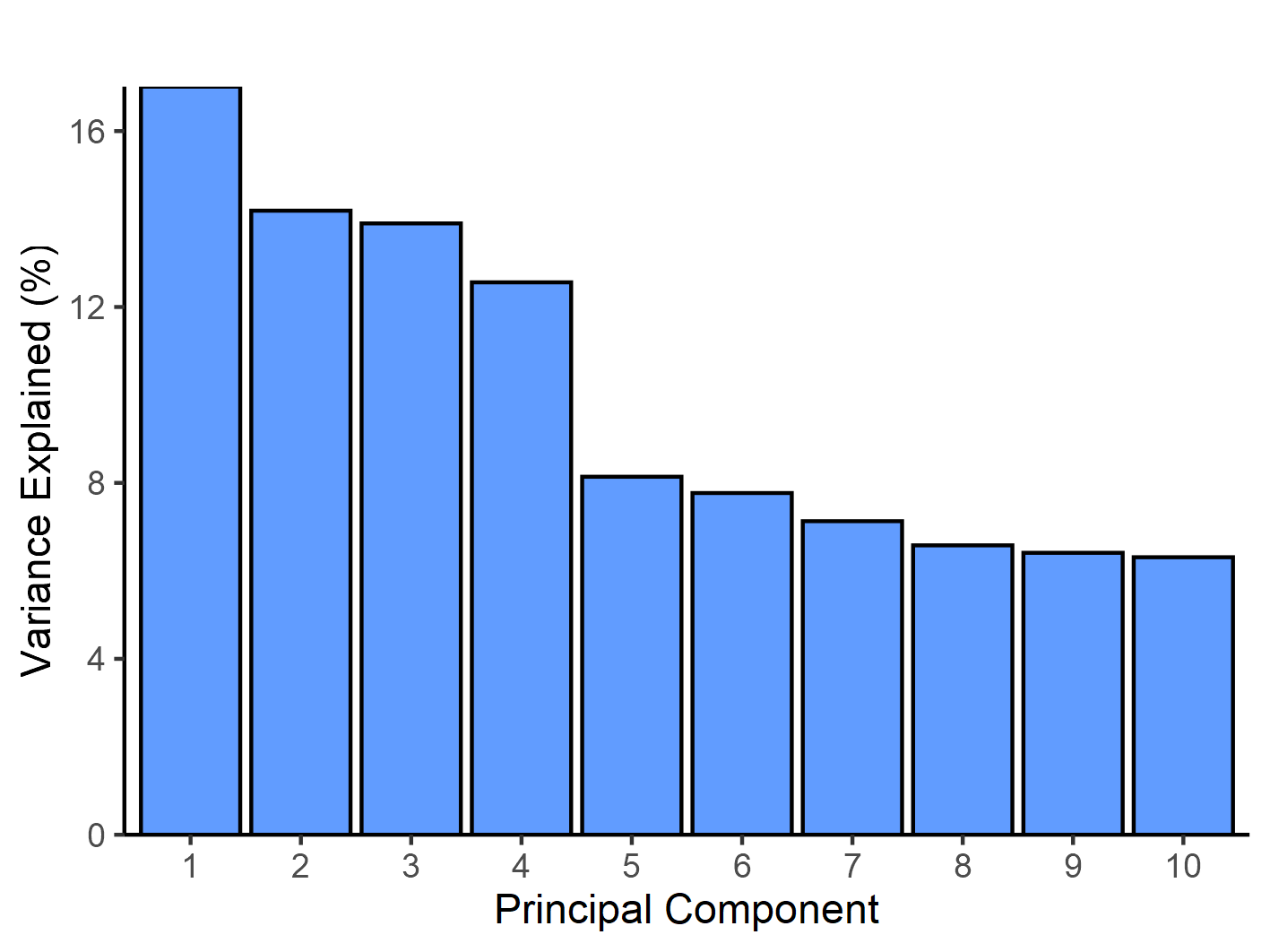

**Supplementary Fig. 4 – Variance in genome-wide SNPs explained by the first 10 principal components in *Quercus robur*.** The plot for PC1 and PC2 is available in the main text (Figure 2c).

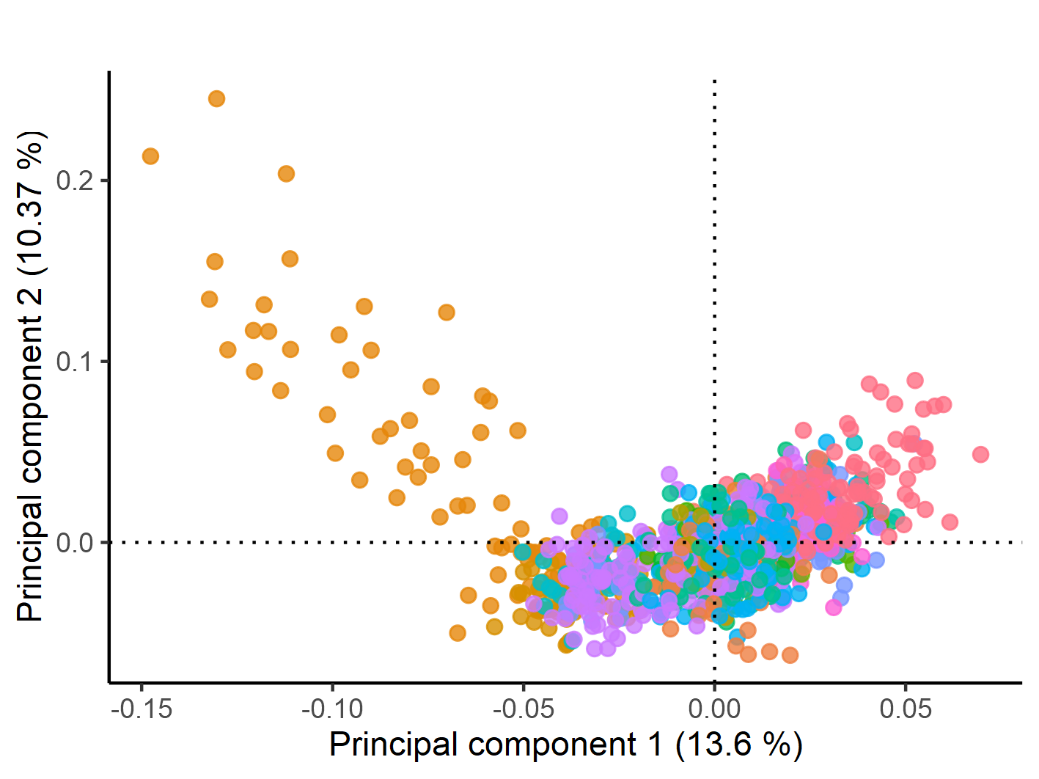

**Supplementary Fig. 5 – Principal components of *Quercus robur* populations after filtering for relatedness up to third degree.** Points are coloured by the collection site. Despite filtering, some individuals from Blickling (brown) are still separated on the PC1 and PC2 axes.

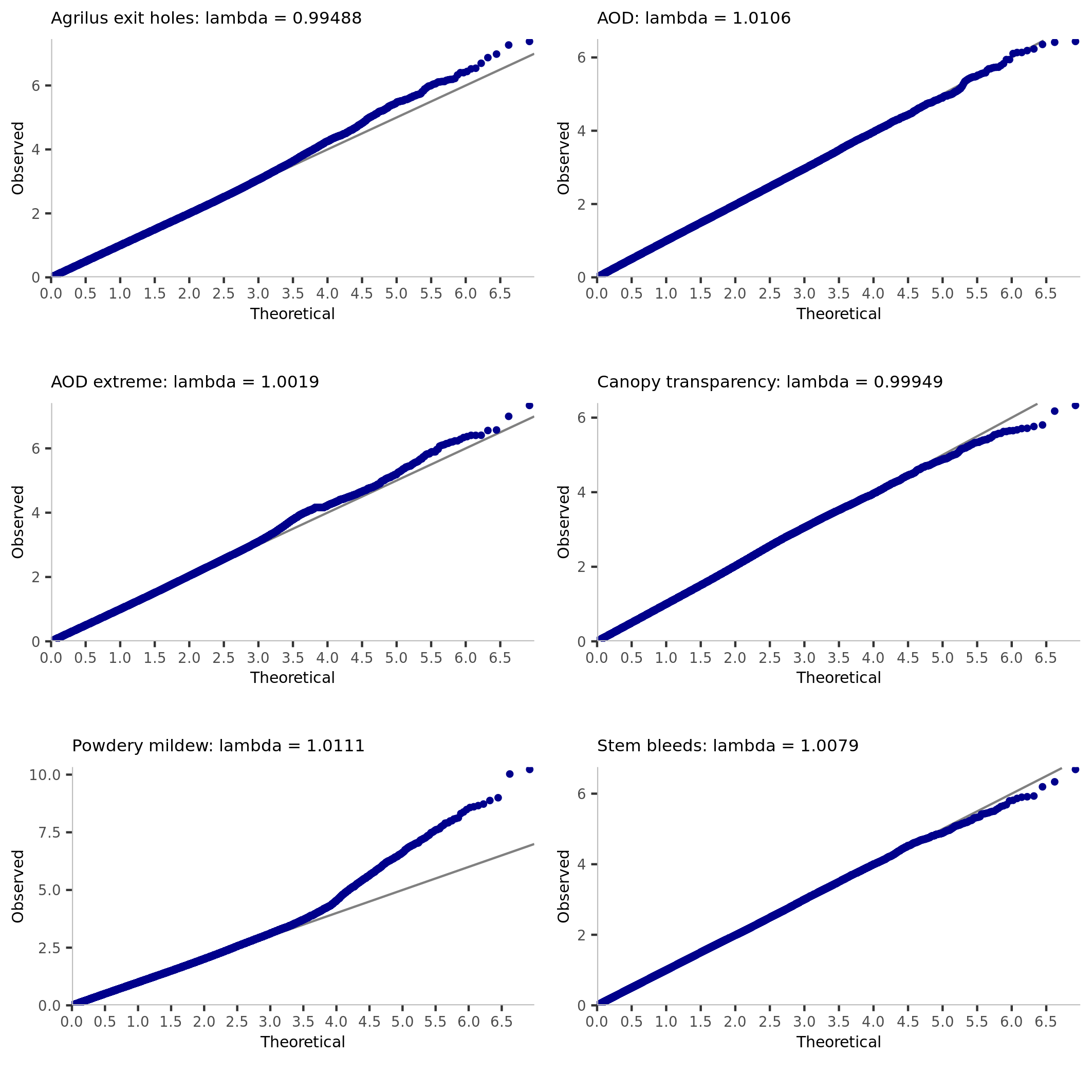

**Supplementary Figure 6 - Quantile-Quantile (QQ) plots of GWAS results for multiple traits in pedunculate oak (*Quercus robur*).** Manhattan plots associated with these results are provided in Figure 4.

**SUPPLEMENTARY TABLES**

| Supplementary Table 1 – Summary statistics for species occurrence and number of *Agrilus,* AOD, and mildew-affected individuals by site. | | | | | | | | | |
| --- | --- | --- | --- | --- | --- | --- | --- | --- | --- |
| Site | Lat | Lng | Sample size | *Quercus petraea* | Hybrid | *Quercus robur* | AOD* | Mildew* | *Agrilus** |
| Abbeydale | 53.33 | -1.51 | 8 | 8 | 0 | 0 | 0 | 0 | 0 |
| Ampthill | 52.04 | -0.45 | 8 | 0 | 0 | 8 | 0 | 0 | 0 |
| Ancaster | 52.75 | -0.48 | 9 | 0 | 1 | 8 | 0 | 0 | 0 |
| Attingham | 52.69 | -2.67 | 101 | 12 | 13 | 76 | 25 | 10 | 15 |
| Big Wood | 52.30 | 1.57 | 11 | 0 | 0 | 11 | 3 | 0 | 1 |
| Blickling | 52.81 | 1.22 | 174 | 0 | 14 | 160 | 85 | 2 | 30 |
| Bryn Engan | 53.10 | -3.90 | 10 | 7 | 3 | 0 | 0 | 0 | 0 |
| Buckminster | 52.81 | -0.71 | 6 | 0 | 0 | 6 | 0 | 0 | 0 |
| Bushy Park | 51.41 | -0.33 | 68 | 0 | 7 | 61 | 34 | 7 | 22 |
| Cannop | 51.81 | -2.56 | 6 | 1 | 0 | 5 | 0 | 0 | 1 |
| Castle Howard Brandrith wood | 54.11 | -0.92 | 6 | 0 | 3 | 3 | 0 | 0 | 0 |
| Castle Howard Cum Hagg wood | 54.13 | -0.94 | 10 | 3 | 5 | 2 | 0 | 0 | 0 |
| Chepstow Park | 51.68 | -2.73 | 10 | 8 | 0 | 2 | 0 | 0 | 0 |
| Chestnuts | 51.83 | -2.47 | 26 | 13 | 2 | 11 | 0 | 0 | 0 |
| Coed Penrhyn | 52.55 | -3.95 | 12 | 5 | 6 | 1 | 0 | 2 | 0 |
| Coed Sarnau | 52.31 | -3.53 | 10 | 7 | 3 | 0 | 0 | 1 | 0 |
| Coedglanllyn | 52.87 | -3.66 | 8 | 3 | 2 | 3 | 0 | 0 | 0 |
| Craigellachie | 57.43 | -3.37 | 10 | 0 | 10 | 0 | 0 | 0 | 0 |
| Crompton | 54.32 | -0.76 | 9 | 4 | 5 | 0 | 0 | 0 | 0 |
| Crowthers Coppice | 52.69 | -3.12 | 17 | 17 | 0 | 0 | 0 | 0 | 0 |
| Delamere | 53.25 | -2.70 | 9 | 3 | 1 | 5 | 0 | 0 | 0 |
| Devichoys Wood | 50.20 | -5.12 | 6 | 6 | 0 | 0 | 0 | 0 | 0 |
| Eastnor Castle | 52.03 | -2.37 | 11 | 0 | 1 | 10 | 4 | 0 | 0 |
| Emilys wood | 52.47 | 0.64 | 7 | 0 | 0 | 7 | 1 | 0 | 0 |
| Epping Site | 51.71 | 0.12 | 7 | 0 | 0 | 7 | 1 | 0 | 0 |
| Fanny Grove | 53.23 | -1.08 | 9 | 5 | 1 | 3 | 1 | 0 | 0 |
| Foxhunting Inclosure | 50.84 | -1.45 | 5 | 0 | 0 | 5 | 0 | 1 | 0 |
| Gillfield Woods | 51.61 | -0.76 | 4 | 0 | 0 | 4 | 0 | 0 | 0 |
| Glen Moriston | 57.21 | -4.61 | 5 | 0 | 5 | 0 | 0 | 0 | 0 |
| Glen Tig | 55.11 | -4.93 | 6 | 6 | 0 | 0 | 0 | 0 | 0 |
| Goose Green | 51.16 | -0.85 | 7 | 0 | 0 | 7 | 0 | 0 | 0 |
| Great Monks | 51.90 | 0.64 | 53 | 2 | 2 | 49 | 37 | 0 | 3 |
| Harewood hall | 53.89 | -1.54 | 7 | 0 | 1 | 6 | 0 | 0 | 0 |
| Hatchlands | 51.26 | -0.47 | 91 | 4 | 7 | 80 | 29 | 4 | 8 |
| Hatfield House | 51.75 | -0.21 | 5 | 3 | 2 | 0 | 1 | 0 | 0 |
| Hazelborough | 52.07 | -1.05 | 5 | 0 | 0 | 5 | 0 | 0 | 0 |
| Hearts of England | 52.25 | -1.85 | 20 | 0 | 0 | 20 | 15 | 1 | 1 |
| Hendre | 51.81 | -2.76 | 5 | 5 | 0 | 0 | 0 | 0 | 0 |
| Highclere Wood | 51.33 | -1.37 | 10 | 0 | 1 | 9 | 0 | 0 | 0 |
| Holkham Hall | 52.96 | 0.81 | 20 | 0 | 0 | 20 | 15 | 0 | 6 |
| Ickworth | 52.23 | 0.66 | 36 | 0 | 2 | 34 | 18 | 3 | 13 |
| Island Thorns | 50.94 | -1.69 | 9 | 3 | 0 | 6 | 0 | 0 | 1 |
| Kellas | 57.57 | -3.42 | 13 | 6 | 7 | 0 | 0 | 0 | 0 |
| Kent College | 51.16 | 0.33 | 4 | 0 | 0 | 4 | 2 | 0 | 1 |
| Kesteven | 53.27 | -0.34 | 6 | 0 | 0 | 6 | 0 | 0 | 0 |
| Kew | 51.48 | -0.30 | 1 | 0 | 0 | 1 | 0 | 0 | 0 |
| Langdale | 52.08 | -2.31 | 223 | 1 | 7 | 215 | 52 | 41 | 10 |
| Langdon Hills | 51.54 | 0.44 | 3 | 2 | 0 | 1 | 0 | 0 | 0 |
| Lineage Wood | 52.10 | 0.76 | 5 | 0 | 0 | 5 | 0 | 0 | 0 |
| Mabie | 55.03 | -3.67 | 13 | 1 | 12 | 0 | 0 | 0 | 0 |
| Maldon Wood | 51.83 | 1.11 | 4 | 0 | 0 | 4 | 0 | 0 | 0 |
| Mitterdale | 54.39 | -3.34 | 4 | 2 | 2 | 0 | 0 | 0 | 0 |
| Moccas Court | 52.08 | -2.93 | 22 | 1 | 1 | 20 | 7 | 0 | 6 |
| Moccas Park | 52.08 | -2.95 | 88 | 3 | 6 | 79 | 41 | 4 | 17 |
| Newlands | 52.72 | -2.07 | 10 | 0 | 1 | 9 | 0 | 0 | 0 |
| Oakamoor | 53.00 | -1.94 | 10 | 9 | 1 | 0 | 0 | 0 | 0 |
| Orlestone Wood | 51.09 | 0.83 | 4 | 0 | 0 | 4 | 1 | 0 | 0 |
| Plora Wood | 51.44 | -0.27 | 6 | 0 | 6 | 0 | 0 | 1 | 0 |
| Red Lodge Wood | 55.62 | -3.03 | 9 | 3 | 5 | 1 | 0 | 0 | 0 |
| Richmond | 51.60 | -1.92 | 264 | 1 | 8 | 255 | 99 | 10 | 61 |
| Rivers Wood | 51.03 | -0.11 | 11 | 0 | 1 | 10 | 0 | 0 | 0 |
| Salcey Forest | 52.15 | -0.83 | 7 | 0 | 0 | 7 | 0 | 0 | 0 |
| Sandringham | 52.82 | 0.50 | 8 | 0 | 0 | 8 | 1 | 0 | 0 |
| Scale green | 54.33 | -3.03 | 3 | 2 | 1 | 0 | 0 | 0 | 0 |
| Scotts Wood | 52.37 | -2.78 | 5 | 3 | 2 | 0 | 0 | 0 | 0 |
| Sherrards Wood | 51.81 | -0.21 | 49 | 24 | 8 | 17 | 23 | 0 | 8 |
| South Forest | 51.43 | -0.65 | 9 | 0 | 0 | 9 | 2 | 0 | 0 |
| Southwick | 52.52 | -0.53 | 45 | 0 | 0 | 45 | 15 | 0 | 0 |
| Spinningdale | 57.87 | -4.24 | 5 | 0 | 5 | 0 | 0 | 0 | 0 |
| Sutton Coldfield | 52.58 | -1.85 | 8 | 2 | 5 | 1 | 0 | 0 | 0 |
| Swaffham | 52.65 | 0.61 | 7 | 0 | 1 | 6 | 0 | 0 | 0 |
| The Straits | 51.16 | -0.86 | 9 | 0 | 0 | 9 | 0 | 0 | 0 |
| Torrachilty | 57.58 | -4.62 | 9 | 1 | 8 | 0 | 0 | 0 | 0 |
| Weald Country | 51.63 | 0.27 | 6 | 0 | 0 | 6 | 0 | 0 | 1 |
| Wensum | 52.71 | 1.26 | 8 | 0 | 1 | 7 | 0 | 0 | 0 |
| Whits Wood | 51.17 | -3.46 | 5 | 5 | 0 | 0 | 0 | 1 | 0 |
| Winding Wood | 52.05 | 0.84 | 123 | 0 | 1 | 122 | 47 | 0 | 7 |
| Wytham Wood | 51.78 | -1.34 | 6 | 0 | 0 | 6 | 2 | 0 | 0 |
| Note: AOD = acute oak decline; Lat = latitude; Lng = longitude; * number of trees affected. | | | | | | | | | |

| Supplementary Table 2 – Environmental variables used as covariates in genome-wide association studies by site. | | | | | | |
| --- | --- | --- | --- | --- | --- | --- |
| Site | Rainfall | Elevation | Ca/Mg | Total SO^2^/SO^4^ | Total NO^2^/NO^3^ | DDEG |
| Abbeydale | 839.32 | 134.98 | 8.7 | 3.2 | 14.4 | 518.51 |
| Ampthill | 637.27 | 99.42 | 4.1 | 2.9 | 16.7 | 526.67 |
| Ancaster | 628.37 | 58.43 | 4.6 | 2.9 | 14.6 | 521.74 |
| Attingham | 681.58 | 54.18 | 2.4 | 1.6 | 9.2 | 509.34 |
| Big Wood | 638.67 | 21.63 | 4.4 | 2.5 | 13.1 | 570.10 |
| Blickling | 736.86 | 35.70 | 4.9 | 2.5 | 12.3 | 525.66 |
| Bryn Engan | 2321.18 | 178.02 | 5.3 | 2 | 8 | 288.51 |
| Buckminster | 699.73 | 119.67 | 5.1 | 3.1 | 16.5 | 465.73 |
| Bushy Park | 620.36 | 8.98 | 5.1 | 2.5 | 14.3 | 778.86 |
| Cannop | 935.88 | 93.82 | 3.9 | 1.9 | 10.7 | 496.45 |
| Castle Howard BRANDRITH WOOD | 721.80 | 68.83 | 4.9 | 2.5 | 9.3 | 403.93 |
| Castle Howard. Cum Hagg Wood | 761.10 | 86.05 | 4.5 | 2.5 | 9.2 | 378.75 |
| Chepstow Park | 1167.48 | 268.15 | 3.4 | 1.9 | 10.6 | 433.33 |
| Chestnuts | 957.74 | 182.70 | 3.9 | 1.8 | 10.3 | 464.97 |
| Coed Penrhyn | 1348.05 | 35.47 | 1.2 | 1.5 | 5.7 | 516.42 |
| Coed Sarnau | 1447.43 | 306.53 | 3.4 | 1.6 | 8.6 | 220.77 |
| Coedglanllyn | 1770.25 | 229.83 | 4 | 1.7 | 7.6 | 296.84 |
| Craigellachie | 931.89 | 182.07 | 1.3 | 0.7 | 3.9 | 122.81 |
| Crompton | 862.15 | 141.40 | 4.6 | 2.6 | 10.6 | 315.56 |
| Crowthers Coppice | 927.83 | 149.25 | 2.2 | 1.4 | 7.5 | 396.04 |
| Delamere | 805.98 | 109.15 | 2.7 | 1.9 | 9.5 | 403.07 |
| Devichoys Wood | 1247.67 | 19.47 | 2.2 | 1.6 | 7.5 | 531.63 |
| Eastnor Castle | 702.13 | 113.88 | 2.8 | 1.8 | 10.6 | 544.98 |
| Emilys wood | 653.53 | 15.65 | 5 | 2.5 | 12.8 | 513.34 |
| Epping Site | 657.49 | 107.55 | 5 | 2.8 | 16.9 | 597.17 |
| Fanny Grove | 646.87 | 50.32 | 5.3 | 3.1 | 12.4 | 467.38 |
| Foxhunting Inclosure | 933.75 | 15.92 | 4.7 | 2.1 | 10.4 | 669.99 |
| Gillfield Woods | 772.11 | 102.17 | 5.5 | 2.7 | 17.2 | 601.83 |
| Glen Moriston | 1281.37 | 123.70 | 1.9 | 0.7 | 3.1 | 265.64 |
| Glen Tig | 1492.40 | 77.60 | 3.2 | 1.1 | 4.3 | 279.24 |
| Goose Green | 824.96 | 89.52 | 6.6 | 2.5 | 13.9 | 581.99 |
| Great Monks | 617.09 | 68.83 | 4.5 | 2.7 | 15.5 | 634.73 |
| Harewood hall | 678.82 | 57.30 | 4.1 | 2.6 | 10.3 | 424.61 |
| Hatchlands | 756.71 | 82.08 | 6.2 | 2.6 | 14.4 | 577.99 |
| Hatfield House | 689.47 | 109.45 | 4.8 | 2.9 | 17.9 | 570.13 |
| Hazelborough | 705.51 | 137.23 | 4.7 | 2.8 | 16.7 | 460.65 |
| Hearts of England | 736.87 | 73.63 | 3.8 | 2.1 | 13 | 530.66 |
| Hendre | 967.38 | 103.57 | 3.1 | 1.7 | 8.7 | 531.98 |
| Highclere Wood | 909.88 | 155.23 | 6 | 2.5 | 15.6 | 458.71 |
| Holkham Hall | 638.65 | 16.35 | 4.8 | 2.6 | 11.8 | 550.79 |
| Ickworth | 649.48 | 78.02 | 5 | 2.7 | 15.6 | 613.16 |
| Island Thorns | 974.17 | 87.15 | 5.1 | 2.2 | 11.7 | 491.38 |
| Kellas | 958.46 | 169.80 | 1.4 | 0.7 | 4 | 209.02 |
| Kent College | 758.01 | 64.10 | 7.7 | 2.6 | 14.1 | 588.78 |
| Kesteven | 646.41 | 16.17 | 4.2 | 3 | 12.2 | 511.44 |
| Kew | 626.16 | 6.65 | 4.9 | 2.5 | 15.2 | 753.32 |
| Langdale | 713.39 | 49.88 | 3.1 | 1.8 | 11.1 | 660.61 |
| Langdon Hills | 584.16 | 39.45 | 4.3 | 2.7 | 15.6 | 651.77 |
| Lineage Wood | 622.74 | 73.52 | 4.4 | 2.7 | 15.3 | 570.94 |
| Mabie | 1266.20 | 97.50 | 1.4 | 0.8 | 4.1 | 317.13 |
| Maldon Wood | 568.43 | 25.27 | 3.8 | 2.6 | 14.9 | 663.00 |
| Mitterdale | 1880.08 | 75.25 | 1.5 | 1.3 | 5.8 | 347.50 |
| Moccas Court | 721.92 | 73.73 | 3 | 1.5 | 8.1 | 520.32 |
| Moccas Park | 748.00 | 75.60 | 3.6 | 1.7 | 9.2 | 521.58 |
| Newlands | 725.69 | 119.70 | 4.5 | 2.4 | 14.3 | 462.25 |
| Oakamoor | 952.33 | 164.10 | 7.5 | 2.6 | 14 | 377.07 |
| Orlestone Wood | 751.94 | 43.30 | 6.3 | 2.7 | 15.3 | 642.42 |
| Plora Wood | 1120.10 | 143.35 | 2.6 | 1.6 | 6.4 | 160.48 |
| Red Lodge Wood | 759.08 | 102.87 | 3.6 | 2.1 | 12.6 | 539.22 |
| Richmond | 670.90 | 39.57 | 5.3 | 2.5 | 14.5 | 727.84 |
| Rivers Wood | 903.01 | 53.22 | 7.1 | 2.5 | 12.9 | 637.48 |
| Salcey Forest | 696.02 | 114.65 | 4.3 | 2.8 | 16.6 | 495.82 |
| Sandringham | 686.81 | 39.82 | 4.8 | 2.7 | 13 | 555.35 |
| Scale green | 2039.50 | 166.05 | 2.7 | 1.5 | 7 | 227.70 |
| Scotts Wood | 778.00 | 104.50 | 2.4 | 1.6 | 8.9 | 465.45 |
| Sherrards Wood | 678.88 | 122.38 | 5.1 | 2.9 | 17.9 | 589.28 |
| South Forest | 717.63 | 69.33 | 5 | 2.6 | 14.9 | 586.27 |
| Southwick | 663.13 | 76.20 | 4.6 | 2.8 | 14.8 | 517.98 |
| Spinningdale | 914.29 | 70.72 | 1.2 | 0.5 | 2.4 | 257.57 |
| Sutton Coldfield | 785.42 | 152.35 | 4.3 | 2.4 | 15.3 | 494.04 |
| Swaffham | 709.29 | 33.07 | 5.3 | 2.6 | 12.8 | 592.38 |
| The Straits | 826.88 | 77.90 | 6.7 | 2.5 | 14.2 | 581.02 |
| Torrachilty | 918.59 | 33.23 | 1.4 | 0.5 | 2.3 | 280.45 |
| Weald Country | 612.58 | 99.68 | 4.9 | 2.8 | 16.2 | 596.75 |
| Wensum | 688.71 | 32.15 | 4.8 | 2.5 | 12.3 | 535.57 |
| Whits Wood | 1040.99 | 94.15 | 3.8 | 1.8 | 8.8 | 519.27 |
| Winding Wood | 605.11 | 63.27 | 4.4 | 2.7 | 15.4 | 592.31 |
| Wytham Wood | 669.53 | 68.18 | 3.9 | 2.4 | 14.1 | 634.89 |
| Notes: We used climate data from the MetOffice HADgridUK at 1km scale. We used monthly average temperature to approximate Day Degree values. Deposition data was retrieved from the CEH and at 5km scale. Elevation was retrieved from OS and then averaged to form a 1ha grid. Rainfall was averaged between 1991 to 2000 at 1km scale. DDEG = day degrees. | | | | | | |

| Supplementary Table 3 – SNPs associated with acute oak decline using an arbitrary cutoff (-log10 >= 5). The gene annotation closest to the SNP position, and its putative function, are provided. | | | | | | |
| --- | --- | --- | --- | --- | --- | --- |
| Chromosome | SNP (bp) | -log10(p) | MAF | Distance (bp) | Gene ID | Putative function |
| Chr1 | 27414889 | 5.13 | 0.08 | 9448 | Qrob_T0605000.2 | Aluminum-activated malate transporter 8 |
| Chr1 | 34374793 | 5.11 | 0.18 | 46968 | Qrob_T0322910.2 | Cathepsin B. |
| Chr1 | 38901547 | 5.37 | 0.13 | 1795 | Qrob_T0183890.2 | Methyltransferase |
| Chr2 | 38097675 | 5.13 | 0.08 | 7430 | Qrob_T0227960.2 | Falz-related bromodomain-containing proteins |
| Chr2 | 73062317 | 5.07 | 0.26 | 18852 | Qrob_T0758690.2 | NA |
| Chr2 | 73062410 | 5.00 | 0.26 | 18945 | Qrob_T0758690.2 | NA |
| Chr2 | 73064334 | 5.16 | 0.31 | 20869 | Qrob_T0758690.2 | NA |
| Chr2 | 92013154 | 5.19 | 0.48 | 9846 | Qrob_T0475570.2 | NA |
| Chr2 | 92027421 | 5.48 | 0.28 | 843 | Qrob_T0475560.2 | NA |
| Chr2 | 92027421 | 5.48 | 0.28 | 145 | Qrob_T0475570.2 | NA |
| Chr2 | 102743306 | 6.80 | 0.06 | 4572 | Qrob_T0288370.2 | Nucleosome remodeling factor subunit CAF1/NURF55/MSI1 |
| Chr2 | 102743306 | 6.80 | 0.06 | 0 | Qrob_T0288380.2 | FAD NAD binding oxidoreductases |
| Chr2 | 102743306 | 6.80 | 0.06 | 3209 | Qrob_T0288390.2 | minichromosome maintenance protein 5 (cell division control protein 46) |
| Chr2 | 102748408 | 5.12 | 0.05 | 0 | Qrob_T0288370.2 | Nucleosome remodeling factor subunit CAF1/NURF55/MSI1 |
| Chr2 | 102748408 | 5.12 | 0.05 | 916 | Qrob_T0288380.2 | FAD NAD binding oxidoreductases |
| Chr2 | 102784388 | 5.00 | 0.27 | 0 | Qrob_T0288330.2 | PPR repeat // PPR repeat family |
| Chr2 | 102784388 | 5.00 | 0.27 | 1037 | Qrob_T0288340.2 | Mannosylglycoprotein endo-beta-mannosidase. |
| Chr3 | 13323340 | 5.79 | 0.07 | 7119 | Qrob_T0208740.2 | Molecular chaperone (DnaJ superfamily) |
| Chr3 | 27018495 | 5.30 | 0.46 | 6979 | Qrob_T0721610.2 | Thioredoxin |
| Chr3 | 27018789 | 5.04 | 0.44 | 6685 | Qrob_T0721610.2 | Thioredoxin |
| Chr3 | 27019330 | 5.27 | 0.46 | 6144 | Qrob_T0721610.2 | Thioredoxin |
| Chr3 | 27020086 | 5.22 | 0.44 | 5388 | Qrob_T0721610.2 | Thioredoxin |
| Chr3 | 27023487 | 5.01 | 0.44 | 1987 | Qrob_T0721610.2 | Thioredoxin |
| Chr3 | 27024497 | 5.67 | 0.44 | 977 | Qrob_T0721610.2 | Thioredoxin |
| Chr3 | 27026257 | 5.32 | 0.45 | 0 | Qrob_T0721610.2 | Thioredoxin |
| Chr3 | 27029634 | 6.02 | 0.46 | 0 | Qrob_T0721610.2 | Thioredoxin |
| Chr3 | 27030166 | 5.03 | 0.42 | 0 | Qrob_T0721610.2 | Thioredoxin |
| Chr3 | 27030740 | 5.26 | 0.18 | 0 | Qrob_T0721610.2 | Thioredoxin |
| Chr4 | 11321328 | 5.11 | 0.06 | 25312 | Qrob_T0147140.2 | Bis(5'-nucleosyl)-tetraphosphatase (asymmetrical). |
| Chr4 | 29085443 | 5.69 | 0.12 | 1777 | Qrob_T0500950.2 | PTHR12100 - SEC10 |
| Chr4 | 31980702 | 5.04 | 0.23 | 48414 | Qrob_T0156090.2 | phytochrome-interacting factor 4* |
| Chr4 | 43422035 | 5.16 | 0.20 | 3934 | Qrob_T0593910.2 | Pre-mRNA processing protein PRP39-related |
| Chr5 | 39039300 | 5.63 | 0.41 | 362 | Qrob_T0736160.2 | RNA-directed DNA polymerase. |
| Chr5 | 39039300 | 5.63 | 0.41 | 640 | Qrob_T0736170.2 | telomerase reverse transcriptase |
| Chr5 | 54582486 | 5.08 | 0.10 | 17405 | Qrob_T0260670.2 | NA |
| Chr6 | 6404359 | 5.12 | 0.27 | 1400 | Qrob_T0159630.2 | DNA-directed RNA polymerase III subunit RPC1 |
| Chr6 | 8636118 | 5.38 | 0.31 | 1879 | Qrob_T0701400.2 | Taxadien-5-alpha-ol O-acetyltransferase. |
| Chr6 | 8637040 | 5.05 | 0.30 | 957 | Qrob_T0701400.2 | Taxadien-5-alpha-ol O-acetyltransferase. |
| Chr6 | 8638411 | 5.11 | 0.30 | 0 | Qrob_T0701400.2 | Taxadien-5-alpha-ol O-acetyltransferase. |
| Chr6 | 8676684 | 5.06 | 0.46 | 1796 | Qrob_T0551190.2 | UDP-glucuronate decarboxylase* |
| Chr6 | 9140518 | 5.41 | 0.07 | 1258 | Qrob_T0641370.2 | Ribonuclease P protein subunit P38-related |
| Chr6 | 30306259 | 5.60 | 0.07 | 68115 | Qrob_T0417330.2 | solute carrier family 32 (vesicular inhibitory amino acid transporter) |
| Chr6 | 55242225 | 5.30 | 0.05 | 0 | Qrob_T0238920.2 | Phosphatidylinositol-3,4-bisphosphate 4-phosphatase. |
| Chr7 | 44055150 | 5.60 | 0.06 | 0 | Qrob_T0316950.2 | Threonine ammonia-lyase. |
| Chr8 | 12709529 | 5.10 | 0.10 | 8511 | Qrob_T0397110.2 | WRKY DNA -binding domain |
| Chr8 | 19129447 | 5.18 | 0.10 | 11419 | Qrob_T0034060.2 | Tyrosine kinase |
| Chr8 | 19129452 | 5.18 | 0.10 | 11424 | Qrob_T0034060.2 | Tyrosine kinase |
| Chr8 | 19129459 | 5.18 | 0.10 | 11431 | Qrob_T0034060.2 | Tyrosine kinase |
| Chr8 | 19129460 | 5.18 | 0.10 | 11432 | Qrob_T0034060.2 | Tyrosine kinase |
| Chr8 | 63485618 | 5.66 | 0.29 | 46473 | Qrob_T0398300.2 | Protein of unknown function (DUF674) |
| Chr8 | 64486907 | 5.45 | 0.40 | 8045 | Qrob_T0450730.2 | Heat shock 70kDa protein 1/8 |
| Chr11 | 13839587 | 5.42 | 0.24 | 45012 | Qrob_T0286270.2 | peroxisomal 2,4-dienoyl-CoA reductase |
| Chr11 | 13839835 | 5.66 | 0.24 | 44764 | Qrob_T0286270.2 | peroxisomal 2,4-dienoyl-CoA reductase |
| Chr11 | 13840791 | 5.54 | 0.24 | 43808 | Qrob_T0286270.2 | peroxisomal 2,4-dienoyl-CoA reductase |
| Chr11 | 13841604 | 5.47 | 0.24 | 42995 | Qrob_T0286270.2 | peroxisomal 2,4-dienoyl-CoA reductase |
| Chr11 | 13841888 | 5.30 | 0.23 | 42711 | Qrob_T0286270.2 | peroxisomal 2,4-dienoyl-CoA reductase |
| Chr11 | 13843174 | 5.78 | 0.24 | 41425 | Qrob_T0286270.2 | peroxisomal 2,4-dienoyl-CoA reductase |
| Chr11 | 13844290 | 6.06 | 0.22 | 40309 | Qrob_T0286270.2 | peroxisomal 2,4-dienoyl-CoA reductase |
| Chr11 | 13844555 | 5.42 | 0.23 | 40044 | Qrob_T0286270.2 | peroxisomal 2,4-dienoyl-CoA reductase |
| Chr11 | 13844644 | 5.01 | 0.25 | 39955 | Qrob_T0286270.2 | peroxisomal 2,4-dienoyl-CoA reductase |
| Chr11 | 13846410 | 6.29 | 0.22 | 38189 | Qrob_T0286270.2 | peroxisomal 2,4-dienoyl-CoA reductase |
| Chr11 | 13846558 | 5.22 | 0.23 | 38041 | Qrob_T0286270.2 | peroxisomal 2,4-dienoyl-CoA reductase |
| Chr11 | 13846644 | 5.30 | 0.24 | 37955 | Qrob_T0286270.2 | peroxisomal 2,4-dienoyl-CoA reductase |
| Chr11 | 13848900 | 5.60 | 0.24 | 35699 | Qrob_T0286270.2 | peroxisomal 2,4-dienoyl-CoA reductase |
| Chr11 | 13848923 | 5.96 | 0.24 | 35676 | Qrob_T0286270.2 | peroxisomal 2,4-dienoyl-CoA reductase |
| Chr11 | 13849898 | 5.92 | 0.20 | 34701 | Qrob_T0286270.2 | peroxisomal 2,4-dienoyl-CoA reductase |
| Chr11 | 13850321 | 5.52 | 0.26 | 34278 | Qrob_T0286270.2 | peroxisomal 2,4-dienoyl-CoA reductase |
| Chr11 | 13851665 | 6.06 | 0.22 | 32934 | Qrob_T0286270.2 | peroxisomal 2,4-dienoyl-CoA reductase |
| Chr11 | 13852285 | 5.37 | 0.24 | 32314 | Qrob_T0286270.2 | peroxisomal 2,4-dienoyl-CoA reductase |
| Chr11 | 13853608 | 6.30 | 0.23 | 30991 | Qrob_T0286270.2 | peroxisomal 2,4-dienoyl-CoA reductase |
| Chr11 | 13853695 | 5.41 | 0.23 | 30904 | Qrob_T0286270.2 | peroxisomal 2,4-dienoyl-CoA reductase |
| Chr11 | 13854990 | 5.48 | 0.23 | 29609 | Qrob_T0286270.2 | peroxisomal 2,4-dienoyl-CoA reductase |
| Chr11 | 13854994 | 5.39 | 0.23 | 29605 | Qrob_T0286270.2 | peroxisomal 2,4-dienoyl-CoA reductase |
| Chr11 | 20239537 | 5.26 | 0.16 | 39455 | Qrob_T0075370.2 | NA |
| Chr11 | 20250297 | 5.04 | 0.16 | 50215 | Qrob_T0075370.2 | NA |
| Qrob_H2.3_Sc0000936 | 23361 | 5.20 | 0.13 | 7881 | Qrob_T0287020.2 | Thioesterase superfamily member |
| Qrob_H2.3_Sc0000936 | 23361 | 5.20 | 0.13 | 7187 | Qrob_T0287030.2 | Thioesterase superfamily member |
| Note: BP = base pair; MAF = minor allele frequency. Distance refers to base pair distance from the SNP to the closest gene; * = located in regions highly differentiated between *Quercus petraea* and *Q. robur* (Gathercole et al., 2026) | | | | | | |

| Supplementary Table 4 – SNPs significantly associated with mildew infection. The gene annotation closest to the SNP position, and its putative function, are provided. When more than one SNP is near a gene, we provide statistics for the SNP with the highest -log(p) value. | | | | | | |
| --- | --- | --- | --- | --- | --- | --- |
| Chr | **SNP (bp)** | **-log_10_(p)** | **MAF** | **N_snps_** | **Gene ID** | **Putative function** |
| 2 | 58465488 | 8.38 | 0.11 | 20 | Qrob_T0327210.2 | Peptide-methionine (R)-S-oxide reductase. |
| 2 | 58509625 | 10.23 | 0.12 | 30 | Qrob_T0633960.2 | SNF2 family DNA-dependent ATPase domain-containing protein [Transcription]. |
| 3 | 51395456 | 6.18 | 0.06 | 1 | Qrob_T0030180.2 | F-box associated domain |
| 3 | 51395456 | 6.18 | 0.06 | 1 | Qrob_T0030190.2 | Domain of unknown function (DUF2828) |
| 3 | 49517805 | 5.96 | 0.09 | 1 | Qrob_T0170360.2 | MYB-LIKE DNA-BINDING PROTEIN MYB // SUBFAMILY NOT NAMED |
| 3 | 12310791 | 6.28 | 0.10 | 1 | Qrob_T0360270.2 | NA |
| 3 | 7987008 | 6.18 | 0.14 | 1 | Qrob_T0573110.2 | Plant protein of unknown function |
| 3 | 10615801 | 6.59 | 0.24 | 1 | Qrob_T0683480.2 | Dosage compensation complex subunit MLE [Transcription]. // Copper chaperone [Inorganic ion transport and metabolism]. |
| 4 | 17998863 | 8.72 | 0.12 | 4 | Qrob_T0410690.2 | ENDOGLUCANASE 12-RELATED |
| 6 | 11094880 | 6.66 | 0.07 | 1 | Qrob_T0184630.2 | NA |
| 6 | 11094880 | 6.66 | 0.07 | 1 | Qrob_T0184640.2 | Pathogenesis-related protein Bet v I family |
| 6 | 27200611 | 6.08 | 0.11 | 1 | Qrob_T0256040.2 | NA |
| 6 | 18856343 | 6.89 | 0.07 | 1 | Qrob_T0476830.2 | AMMONIUM TRANSPORTER |
| 6 | 23234007 | 7.05 | 0.07 | 1 | Qrob_T0601940.2 | Dihydroxy-acid dehydratase. |
| 6 | 25097911 | 6.14 | 0.16 | 1 | Qrob_T0678250.2 | Histone H3 (Lys9) methyltransferase SUV39H1/Clr4 |
| 6 | 25039929 | 6.11 | 0.11 | 1 | Qrob_T0678300.2 | NA |
| 6 | 25024910 | 6.10 | 0.11 | 1 | Qrob_T0678310.2 | EPIDIDYMAL MEMBRANE PROTEIN E9-RELATED // SUBFAMILY NOT NAMED |
| 6 | 27279675 | 6.76 | 0.11 | 5 | Qrob_T0690030.2 | PPR repeat // PPR repeat family // Pentatricopeptide repeat domain |
| 6 | 27265217 | 6.15 | 0.11 | 1 | Qrob_T0690040.2 | PPR repeat // PPR repeat // PPR repeat family // Pentatricopeptide repeat domain |
| 6 | 27217418 | 6.91 | 0.16 | 1 | Qrob_T0690090.2 | CYSTEINE-RICH REPEAT SECRETORY PROTEIN 11-RELATED |
| 7 | 41400821 | 6.01 | 0.06 | 2 | Qrob_T0090390.2 | 1-(5-phosphoribosyl)-5- ((5-phosphoribosylamino)methylideneamino)imidazole-4-carboxamide isomerase. |
| 7 | 41400821 | 6.01 | 0.06 | 2 | Qrob_T0090400.2 | HEME-BINDING PROTEIN-RELATED // SUBFAMILY NOT NAMED (PTHR11220:SF22) |
| 7 | 41400821 | 6.01 | 0.06 | 2 | Qrob_T0090410.2 | THIOREDOXIN FAMILY PROTEIN (PTHR11260:SF136) |
| 7 | 1880156 | 6.43 | 0.11 | 3 | Qrob_T0277850.2 | K(+) EFFLUX ANTIPORTER 3, CHLOROPLASTIC (PTHR16254:SF6) |
| 7 | 49125549 | 7.04 | 0.09 | 1 | Qrob_T0647350.2 | PHOSPHATIDYLINOSITOL N-ACETYGLUCOSAMINLYTRANSFERASE SUBUNIT P-LIKE PROTEIN* |
| 8 | 53351942 | 6.31 | 0.06 | 4 | Qrob_T0705670.2 | INACTIVE PURPLE ACID PHOSPHATASE 28-RELATED* |
| 8 | 53352397 | 6.25 | 0.06 | 1 | Qrob_T0705680.2 | Endo-1,4-beta-xylanase. |
| 11 | 29836479 | 6.29 | 0.07 | 1 | Qrob_T0171080.2 | TRANSMEMBRANE PROTEIN HTP-1 RELATED |
| 11 | 47199232 | 6.33 | 0.06 | 1 | Qrob_T0206100.2 | PECTINESTERASE |
| 11 | 47199232 | 6.33 | 0.06 | 1 | Qrob_T0206110.2 | ZINC FINGER FYVE DOMAIN CONTAINING PROTEIN // SUBFAMILY NOT NAMED (PTHR22835:SF150) |
| 11 | 19053978 | 6.36 | 0.14 | 4 | Qrob_T0289900.2 | alcohol dehydrogenase class-P [EC:1.1.1.1] |
| 11 | 19120896 | 6.49 | 0.10 | 3 | Qrob_T0289910.2 | Alcohol dehydrogenase. |
| 11 | 19140930 | 7.99 | 0.17 | 5 | Qrob_T0289920.2 | UDP-glucose 6-dehydrogenase* |
| 11 | 19172270 | 6.06 | 0.17 | 1 | Qrob_T0289930.2 | NA |
| 11 | 19219480 | 6.44 | 0.22 | 1 | Qrob_T0289970.2 | Structural maintenance of chromosome protein 1 (sister chromatid cohesion complex Cohesin<a0> subunit SMC1) |
| 11 | 19233485 | 6.76 | 0.12 | 7 | Qrob_T0289980.2 | leucine-rich PPR motif-containing protein, mitochondrial |
| 11 | 19239887 | 7.23 | 0.09 | 4 | Qrob_T0290010.2 | MYC // SUBFAMILY NOT NAMED |
| 11 | 19274814 | 7.00 | 0.10 | 4 | Qrob_T0290020.2 | L-galactose 1-dehydrogenase. |
| 11 | 19274814 | 7.00 | 0.10 | 2 | Qrob_T0290030.2 | REPLICATION FACTOR A 1, RFA1 // SUBFAMILY NOT NAMED |
| 11 | 19361366 | 7.17 | 0.16 | 6 | Qrob_T0290060.2 | Leucine Rich Repeat // NB-ARC domain // TIR domain |
| 11 | 19381254 | 7.19 | 0.11 | 14 | Qrob_T0290090.2 | Domain of unknown function (DUF313) |
| 11 | 19397281 | 8.66 | 0.16 | 21 | Qrob_T0290100.2 | Glutamate-1-semialdehyde 2,1-aminomutase* |
| 11 | 19421443 | 6.52 | 0.14 | 4 | Qrob_T0290110.2 | METALLOPHOSPHOESTERASE DOMAIN-CONTAINING PROTEIN (PTHR22953:SF18) |
| 11 | 19426413 | 6.78 | 0.14 | 21 | Qrob_T0290120.2 | NA |
| 11 | 24772933 | 7.81 | 0.10 | 5 | Qrob_T0540440.2 | NA |
| 11 | 24772933 | 7.81 | 0.10 | 5 | Qrob_T0540450.2 | Uncharacterized conserved protein [Function unknown]. |
| 11 | 24882696 | 8.00 | 0.10 | 4 | Qrob_T0540470.2 | INORGANIC PYROPHOSPHATASE (PTHR10286:SF3) |
| 12 | 22997373 | 6.42 | 0.08 | 1 | Qrob_T0043980.2 | B-box zinc finger* |
| 12 | 15841352 | 6.30 | 0.16 | 1 | Qrob_T0082210.2 | ZINC FINGER FYVE DOMAIN CONTAINING PROTEIN // SUBFAMILY NOT NAMED (PTHR22835:SF137) |
| Qrob_H2.3_Sc0000249 | 1006056 | 6.64 | 0.11 | 1 | Qrob_T0277900.2 | Tyrosine kinase specific for activated<a0> |
| Qrob_H2.3_Sc0001094 | 36097 | 6.54 | 0.12 | 3 | Qrob_T0306170.2 | NA |
| Qrob_H2.3_Sc0001094 | 36097 | 6.54 | 0.12 | 3 | Qrob_T0306180.2 | F-box domain |
| Notes: MAF = minor allele frequency; N_snps_ = number of significant SNPs near the gene annotation; * = gene located in a highly differentiated genomic region between *Quercus petraea* and *Q. robur* (Gathercole et al., 2026). | | | | | | |
